# Presynaptic inhibition rapidly stabilises recurrent excitation in the face of plasticity

**DOI:** 10.1101/2020.02.11.944082

**Authors:** Laura Bella Naumann, Henning Sprekeler

## Abstract

Hebbian plasticity, a mechanism believed to be the substrate of learning and memory, detects and further enhances correlated neural activity. Because this constitutes an unstable positive feedback loop, it requires additional homeostatic control.

Computational work suggests that in recurrent networks, the homeostatic mechanisms observed in experiments are too slow to compensate instabilities arising from Hebbian plasticity and need to be complemented by rapid compensatory processes. We suggest presynaptic inhibition as a candidate that rapidly provides stability by compensating recurrent excitation induced by Hebbian changes. Presynaptic inhibition is mediated by presynaptic GABA receptors that effectively and reversibly attenuate transmitter release. Activation of these receptors can be triggered by excess network activity, hence providing a stabilising negative feedback loop that weakens recurrent interactions on sub-second timescales. We study the stabilising effect of presynaptic inhibition in a recurrent networks, in which presynaptic inhibition is implemented as a multiplicative reduction of recurrent synaptic weights in response to increasing inhibitory activity. We show that networks with presynaptic inhibition display a gradual increase of firing rates with growing excitatory weights, in contrast to traditional excitatory-inhibitory networks. This alleviates the positive feedback loop between Hebbian plasticity and network activity and thereby allows homeostasis to act on timescales similar to those observed in experiments. Our results generalise to spiking networks with a biophysically more detailed implementation of the presynaptic inhibition mechanism. In conclusion, presynaptic inhibition provides a powerful compensatory mechanism that rapidly reduces effective recurrent interactions and thereby stabilises Hebbian learning.

## Introduction

Synaptic plasticity is widely believed to be the neuronal substrate for learning and memory. Hebbian plasticity [20], in particular, is the long-standing prime candidate for associative learning. It preferentially connects neurons that are co-active, thereby linking neural representations of events that co-occur in time. Notwithstanding its obvious suitability for learning, Hebbian plasticity also has the precarious side-effect of creating a positive feedback loop [2]. Pairs of neurons whose connection was strengthened by Hebbian plasticity tend to be even more co-active, which in turn further strengthens their connection. The standard argument why this vicious circle does not generate runaway activity in the brain is that there is a broad spectrum of homeostatic mechanisms that keep this instability at bay [2, 24, 52]. Such mechanisms have been demonstrated in various forms, including homeostatic scaling [51], intrinsic plasticity [**?**, 15], metaplasticity [6, 26] or plasticity of inhibition [45, 56]. While these mechanisms counteract modifications that take neuronal activity out of a functional regime, they all occur on a relatively long time scale of hours or days. They are therefore poorly suited to protect neural circuits against unwanted consequences of rapid changes in input or connectivity [59, 60]. Zenke et al. [59] recently suggested that the known homeostatic mechanisms must hence be complemented by rapid compensatory processes that render the circuit stable on shorter time scales and hence give homeostasis the time it needs.

We suggest that presynaptic inhibition – a form of inhibition that has attained little attention in computational modelling – could serve as a candidate for such a rapid compensatory process. Presynaptic inhibition is a mechanism that suppresses synaptic transmission by means of presynaptic receptors and can occur through a variety of pathways [30]. At excitatory synapses, activation of presynaptic GABA_B_ receptors causes a reduction of neurotransmitter release [25] by inhibiting voltage-dependent Calcium channels [22, 58]. In turn, presynaptic GABA_B_ receptors are activated by interneuron-mediated GABA release [53], potentially by means of GABA spillover [4, 16, 22]. Hence, excess activity in excitatory neurons can recruit inhibitory interneurons, which in turn activate presynaptic inhibition and thereby suppress recurrent excitatory connections. This provides a dynamic and reversible negative feedback loop onto recurrent excitation – a potent source of network instability – on timescales of hundreds of milliseconds [9, 32, 53]. GABA_B_-mediated presynaptic inhibition has been observed in a range of brain areas of different species (see for example [14, 18, 25, 35, 53]), but the functional implications, in particular in recurrent circuits, remain elusive.

Here, we use simulations of rate-based networks as well as mathematical analyses to study the compensatory properties of presynaptic inhibition in recurrent circuits. We show that presynaptic inhibition ensures a gradual increase of firing rates for increases in recurrent excitation, in contrast to traditional excitatory-inhibitory networks [11]. This stabilises neural activity in networks subject to Hebbian plasticity even if homeostasis is slow. We find that the stabilising properties are robust to network parameters and details of the mechanism. Finally, we show that these results generalise to networks of spiking neurons.

## Results

To study the compensatory effects of presynaptic inhibition, we simulated networks of excitatory and inhibitory rate-based neurons and analysed the effect of changes at excitatory recurrent weights on network activity. We start by systematically increasing recurrent excitation to illustrate the stabilising properties of presynaptic inhibition. Then, we study the interaction of presynaptic inhibition with the Bienenstock-Cooper-Munro (BCM) rule, a rate-based Hebbian plasticity rule that comprises a homeostatic control mechanism. We vary the timescale of homeostatic control and contrast the behaviour of networks with and without presynaptic inhibition.

### Presynaptic inhibition in a recurrent network model

Our model of presynaptic inhibition is based on the following biophysical mechanism: GABA spillover from inhibitory synapses activates presynaptic GABA_B_ receptors at excitatory synapses. This suppresses voltage-dependent calcium (Ca^2+^) channels, thereby inhibiting neurotransmitter release. Thus, activation of inhibitory neurons can lead to presynaptic inhibition of excitatory synaptic transmission [53] (see Fig. 1A for a schematic illustration).

**Figure 1.**
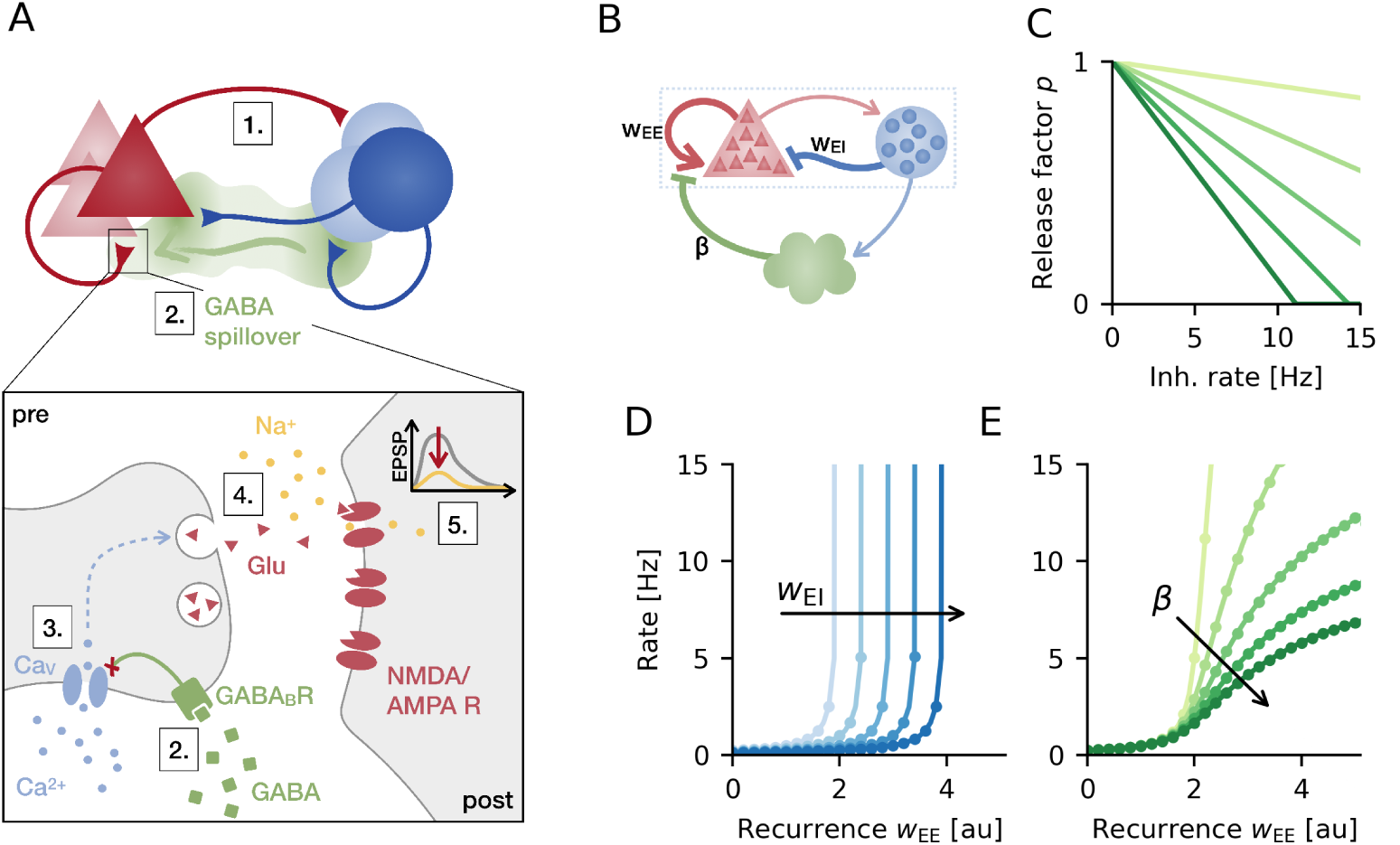
Presynaptic inhibition in a recurrent network. **A.** Presynaptic inhibition mechanism: (1) Network activity drives inhibitory interneurons. (2) GABA released by interneurons can spill over to nearby excitatory synapses, where it binds to presynaptic GABA_B_ receptors that (3) in turn inhibit voltage-sensitive Calcium (Ca^2+^) channels. (4) Reduced Ca^2+^ influx decreases the release of neurotransmitter Glutamate and (5) therefore the amplitude of EPSPs. **B.** Simplified rate-based model consisting of excitatory (red) and inhibitory population (blue) and a presynaptic inhibition mechanism (green). **C.** Release factor as a linear function of inhibitory neuron activity for different slopes *β* = 0.01, 0.03, …, 0.09. **D.** Population rate as a function of excitatory recurrence (*w*_EE_) without presynaptic inhibition for increasing strength of postsynaptic inhibition (weight *w*_EI_ = 1, 1.5, …, 3). **E.** Same as D but for fixed postsynaptic inhibition and increasing strength of presynaptic inhibition (slope of transfer function *β*, see C). Markers are simulation results and solid lines analytically determined steady-state rates (see SI Appendix, Methods and Models).

In the rate network, we use inhibitory activity as a proxy of GABA spillover, which in turn modulates excitatory synaptic transmission (Fig. 1B). We model this presynaptic modulation as a multiplicative “release” factor that scales excitatory synaptic weights and decreases with increasing inhibitory activity. For the sake of analytical tractability, the release factor decreases linearly with inhibitory firing rate (Fig. 1C). Therefore, the sensitivity of presynaptic inhibition to inhibitory firing rate (i.e. the strength of presynaptic inhibition) is determined by the slope of this linear transfer function. Here, we assume that the effect of presynaptic inhibition is homogeneous and diffuse enough such that all inhibitory neurons contribute to the modulation of a global release factor. This global factor then multiplicatively scales all excitatory recurrent weights in the network. Note that the presented results remain unchanged for model variants, in which presynaptic inhibition acts more locally, because we exclusively study random networks without spatial structure.

### Presynaptic inhibition compensates for recurrent excitation

Hebbian plasticity adjusts recurrent synaptic weights according to correlations in neural activity. At the same time, changing recurrent weights affects the activity of interconnected neurons, forming a potentially destabilising positive feedback loop. Thus, how the overall firing rate increases with changes in recurrent excitatory weights is an indicator of stability in the presence of Hebbian plasticity. We therefore first study the effect of ad-hoc homogeneous increases in excitatory recurrence.

In a network without presynaptic inhibition, minor changes in overall excitatory recurrence cause major increases in the mean population firing rate (Fig. 1D). Above a critical value, the recurrent excitation drives the network to pathologically high activity states. Increasing the strength of postsynaptic inhibition does not eliminate the supralinear dependence, but merely shifts it to higher values of the excitatory recurrence. With presynaptic inhibition in place, firing rates have a qualitatively different dependence on recurrence. Network activity gradually increases with excitatory weights for arbitrarily strong recurrent excitation (Fig. 1E). We confirm these results in mathematical analyses and show that the mean population rate saturates at a finite value as recurrent weights increase (SI Appendix, Methods and Models). How much the rate increases with recurrence and where it saturates depends on the strength of presynaptic inhibition (Fig. 1E), which is determined by the transfer function’s slope (Fig. 1C).

In summary, while conventional networks of excitatory and inhibitory populations are prone to instabilities triggered by increases in excitatory recurrence, adding presynaptic inhibition allows for gradual increases of neural activity with growing excitation. This makes presynaptic inhibition a candidate mechanism to break the positive feedback loop generated by Hebbian plasticity and recurrent excitation.

### Presynaptic inhibition prevents runaway excitation in the face of Hebbian plasticity

Does presynaptic inhibition also stabilise Hebbian plasticity at recurrent excitatory synapses? Recent work by Zenke et al. [60] has revealed that stability in spiking networks with Hebbian plasticity on excitatory synapses requires the timescale of homeostasis to be substantially shorter than that of plasticity. To show that presynaptic inhibition increases the range of stability, we first qualitatively reproduce these results in a recurrent rate network with plastic excitatory synapses. To this end, we use the BCM rule [6], which has a correspondence to the triplet rule used in the work of Zenke et al. [19, 38]. Homeostasis is implemented by a sliding threshold in the BCM rule that aims at keeping a running average of single neuron firing rates at a given target rate [6] (Fig. 2A, left). The time constant of this running average determines the timescale of homeostasis and is therefore referred to as the homeostatic time constant in the following [60].

**Figure 2.**
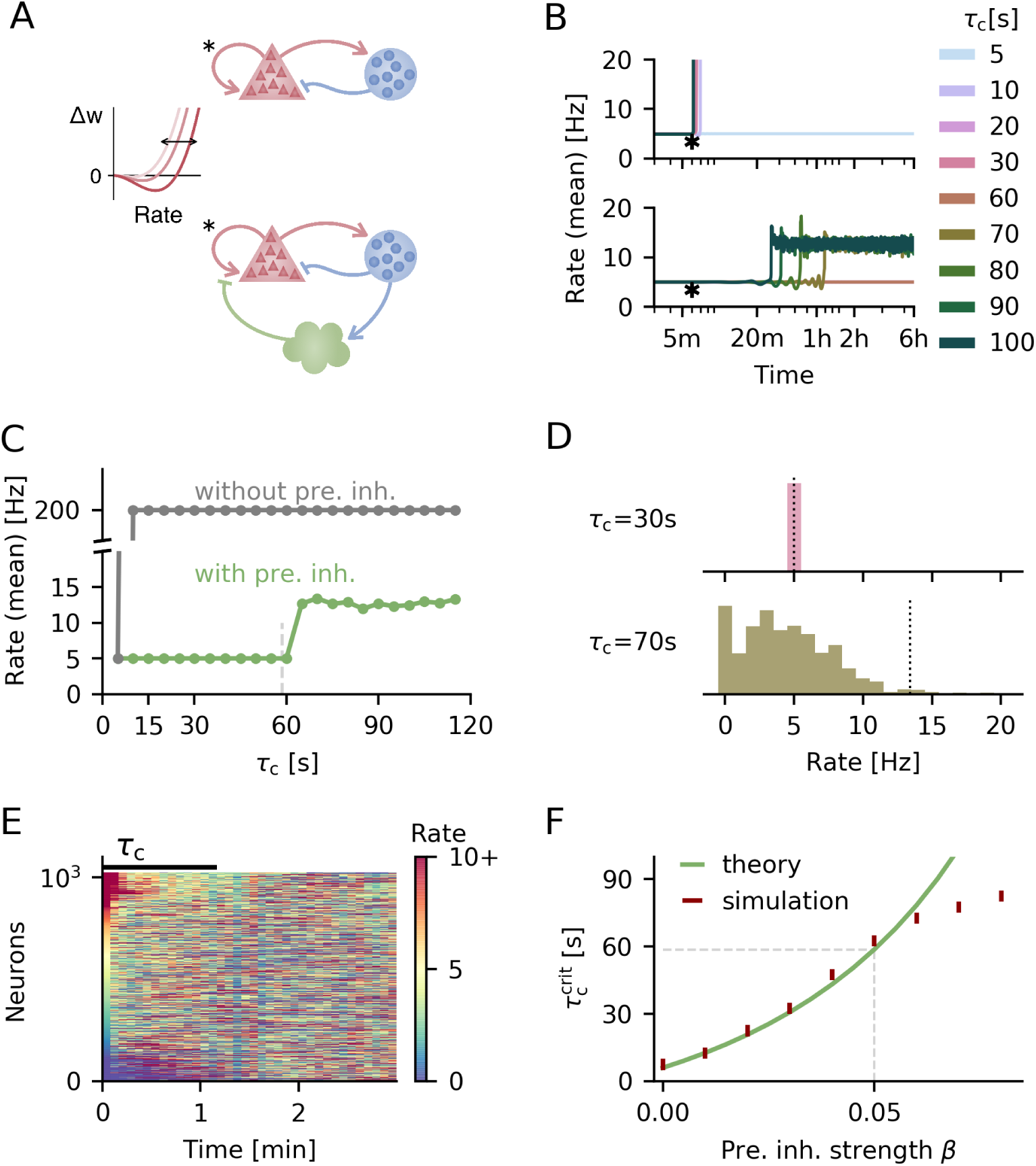
Presynaptic inhibition increases critical homeostatic time constant and maintains population firing rate at low rates. **A.** Circuit diagrams for networks without (top) and with presynaptic inhibition (bottom). Excitatory recurrent synapses are subject to the BCM rule with sliding threshold (illustration on the left). **B.** Temporal evolution of the average firing rate for different homeostatic time constants *τ*_c_ for networks without (top) and with presynaptic inhibition (bottom). Plasticity is switched on after 6 min of simulation time (indicated by asterisk). **C.** Mean final population rate as a function of homeostatic time constant. Vertical dashed line marks critical homeostatic timescale from theory with presynaptic inhibition (see F). **D.** Final firing rate distribution for example runs with presynaptic inhibition for a faster (top) and slower (bottom) homeostatic time constant. Average firing rate is indicated by vertical, dotted line. **E.** Single neuron firing rates over time at the end of the simulation for *τ*_c_ = 70s. Neurons are sorted according to their activity in the beginning of this period. Target rate level (5 Hz) is yellow. **F.** Critical homeostatic time scale as a function of presynaptic inhibition strength *β*. For simulations the vertical bars indicate the range between largest stable and smallest unstable timescale tested. *β* = 0 corresponds to networks without presynaptic inhibition.

In accordance with previous work, we find that as long as homeostasis is fast enough (on the order of a few seconds), the BCM rule keeps network and single neuron activity at the target rate (Fig. 2B, top). However, above a critical homeostatic time constant, the sliding threshold is not able to control the positive feedback loop of recurrent excitation and Hebbian plasticity.

Including presynaptic inhibition in the network (Fig. 2A, bottom) alleviates this instability. We observe two effects: First, the critical homeostatic time constant, below which all neurons remain at the target rate is increased more than tenfold (Fig. 2B, bottom). Second, presynaptic inhibition prevents a pathological runaway of activity even beyond this critical homeostatic timescale (Fig. 2C).

Above the respective critical time constant, the network behaviour is remarkably different between networks with and without presynaptic inhibition. Without it the firing rate jumps from target to the maximum rate even for small homeostatic time constants (Fig. 2C). With presynaptic inhibition, the mean population rate remains low (Fig. 2B, bottom and Fig. 2 C) although single neuron firing rates show a broad distribution of values (Fig. 2D) and large temporal fluctuations (see Fig. 2E, and SI Appendix, Fig. S2 A, B). The low mean firing rate on the population level is the consequence of presynaptic inhibition, which limits the population rate by suppressing recurrent interactions. The broad distribution of firing rates is a result of the BCM rule: neurons firing above the target rate will further increase their activity due to the Hebbian nature of the BCM rule, and the converse applies to neurons firing below the target rate. The slow sliding threshold provides a delayed negative feedback loop to this diversification, leading to a perpetual turnover of those neurons firing at high rates (Fig. 2E) that depends on the homeostatic time constant (SI Appendix, Fig. S2).

To understand what determines the increase in the critical time constant with presynaptic inhibition, we used a similar approach as in previous work [60] (SI Appendix, Methods and Models) to mathematically derive the critical homeostatic time constant with presynaptic inhibition in place (cf. Eq (38)):

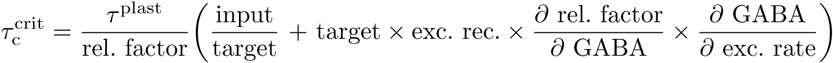

This equation shows that the range of stability increases with

- lower release factor at the target rate,
- stronger excitatory recurrence,
- a steeper decrease in release factor in response to overall GABA spillover,
- a stronger dependence of GABA spillover on excitatory firing rate and
- the amount of background input to the network in relation to the target rate.

The mathematical analysis accurately predicts the critical homeostatic timescale above which firing rates in the network deviate from the target, as long as presynaptic inhibition is not too strong (Fig. 2F). As the mechanism is neuron-unspecific, stronger presynaptic inhibition introduces more competition between neurons in addition to the BCM rule, which can further amplify existing heterogeneities in the network. These heterogeneities are not captured by the mean population model used in the mathematical analysis and thus the critical timescale in the full network can deviate from the theoretical prediction. If we remove the noise on initial excitatory weights – a prominent source of heterogeneity in these networks – the simulation results match the theory regardless of presynaptic inhibition strength (SI Appendix, Fig. S3). Here we focus on the parameter regime in which the theoretical predictions for the critical homeostatic timescale hold despite heterogeneities in the network.

Note that homeostatic timescales need to be considered relative to timescales of plasticity. To limit simulation time, the timescale of plasticity was about one minute in the simulations – shorter than to be expected in biology. Zenke et al. [60] estimated the time scale of plasticity based on slice experiments and obtained an estimate of about 50 minutes. In slice experiments, it is commonly observed that after the induction protocol, synaptic potentials only gradually increase within tens of minutes [43]. With these relative time scales in mind, a critical homeostatic time constant that is similar to the time scale of plasticity (about 60 seconds) is on the order of almost an hour. The increase mediated by presynaptic inhibition may hence render the experimentally observed forms of homeostasis functional.

### Presynaptic inhibition reduces sensitivity to strength of background input

In the model, the BCM rule raises or lowers excitatory recurrent weights to achieve a given target rate. As the background input also contributes to single neuron firing rates, its strength affects the magnitude of recurrent weights necessary to reach the target rate. Because the recurrent weights are critical for the stability of the network, we conjecture that the critical homeostatic timescale is strongly affected by the level of background input. Because presynaptic inhibition provides a negative feedback on the recurrent weights, we expect it to reduce the influence of the background input on network stability.

Indeed, we find that without presynaptic inhibition the range of stability of the network increases strongly with the external input (Fig. 3A, top). For the relatively low background input we used so far, homeostasis needs to be at least ten times faster than the effective timescale of plasticity (5 compared to 60 seconds)(Fig. 3A, top left). Increasing the background input substantially extends the range of stability in networks without presynaptic inhibition (Fig. 3A, top middle and right). If the input is similar to the target rate, homeostasis on the order of the plasticity timescale alone is sufficient to maintain stable firing at the target (see also Fig. 3D).

**Figure 3.**
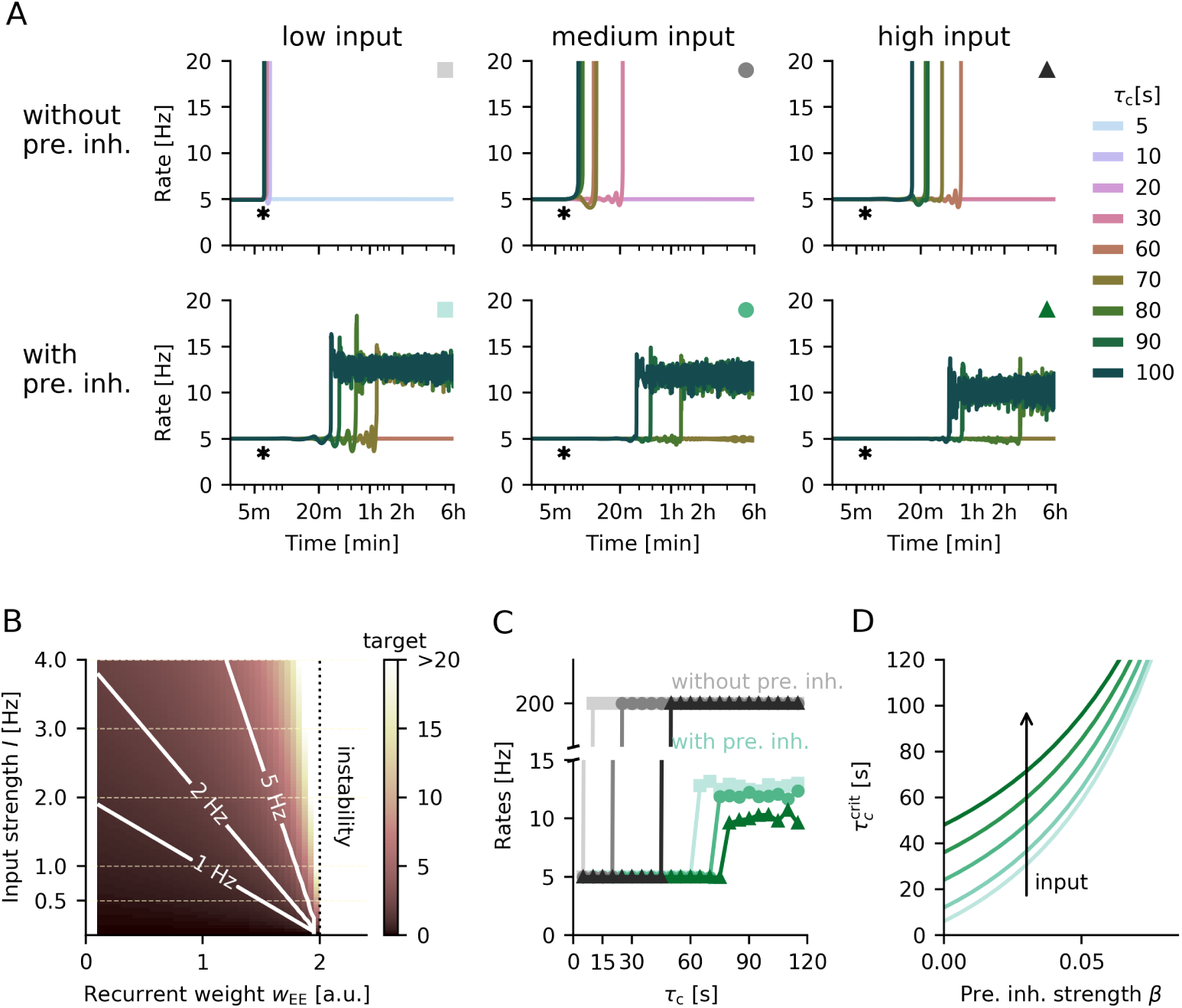
Sensitivity of networks with and without presynaptic mechanism to levels of external input. **A.** Temporal evolution of the average firing rate for different homeostatic time constants *τ*_c_ for networks without (top) or with presynaptic inhibition (bottom) and different levels of background input strength (low - 0.5 Hz, medium - 1 Hz and high - 2.5 Hz). Strength of presynaptic inhibition is *β* = 0.05. **B.** Target firing rate in Hz as a function of external input strength compared to mean recurrent excitatory weight in a network without plasticity and presynaptic inhibition. **C.** Mean population rate as a function of homeostatic time constant in networks with (blue-green) and without (grey) presynaptic inhibition for different strengths of background input. Markers and their colours correspond to A indicating the parametrisation (low, medium or high, respectively). **D.** Critical homeostatic timescale derived analytically as a function of presynaptic inhibition strength for increasing strengths of background input (*I* = 0.5, 1, 2, 3, 5)

The (in)stability of networks without presynaptic inhibition can be understood by revisiting the dependence of output firing rate on recurrent excitatory weights (cf. Fig. 1B). For lower background input, the mean recurrent weight has to be increased closer to the point of instability for the neurons in the population to reach a given target rate (Fig. 3B). In consequence, the network is more prone to destabilisation by the Hebbian contribution of the BCM rule, because small changes in excitatory recurrent weights can push the network over this stability threshold if homeostatic control is too slow.

Presynaptic inhibition removes the sudden point of instability, and introduces a gradual increase of population activity with increasing recurrent excitation (cf. Fig. 1C). In consequence, the range of stability in networks with presynaptic inhibition is less dependent on the strength of background input compared to target rate (Fig. 3A, bottom). Although higher inputs increase the critical homeostatic timescale, this increase is less prominent than in networks without presynaptic inhibition (Fig. 3C). This observation is in line with the analytically determined critical homeostatic timescale: The stronger presynaptic inhibition, the smaller is the difference in critical timescales when varying the strength of background input (Fig. 3D). In addition, when homeostasis is slower than the critical timescale, the mean population rate decreases with higher inputs (Fig. 3C). As higher inputs require weaker recurrence (Fig. 3B), plasticity-induced fluctuations of single neuron firing rates are smaller, leading to a lower mean population rate.

In summary, the critical homeostatic time constant and, accordingly, network stability crucially depend on the relation of background input to the target rate. Weak inputs require compensation by strong recurrent excitation, bringing network dynamics closer to instability. As networks with presynaptic inhibition do not suffer from such instabilities, they have a weaker dependence on the level of background input or excitatory recurrence.

### Presynaptic inhibition in a spiking network confirms results from rate-based model

So far we have considered a strongly simplified linear rate-based neuron model. Moreover, we incorporated GABA spillover indirectly through the inhibitory firing rate. In the following we validate our results in a more complex, spiking network model. To this end, we simulate networks of randomly connected leaky integrate-and-fire neurons with current-based synapses. We include GABA spillover more explicitly: spikes from a given inhibitory neuron cause a local increase in GABA concentration around those excitatory neurons it targets. The accumulated GABA then inhibits afferent excitatory transmission, modelled again as a multiplicative factor onto recurrent synaptic weights (Fig. 4A). Thus higher GABA concentrations leads to a decrease in amplitude of excitatory postsynaptic potentials (EPSP) (Fig. 4B). To simplify a comparison with the rate-based analysis, we rescale the GABA concentration to values comparable to inhibitory or excitatory firing rates in the system (see SI Appendix, Methods and Models) and use the same linear transfer function linking GABA concentration to release factor as before (Fig. 4C).

**Figure 4.**
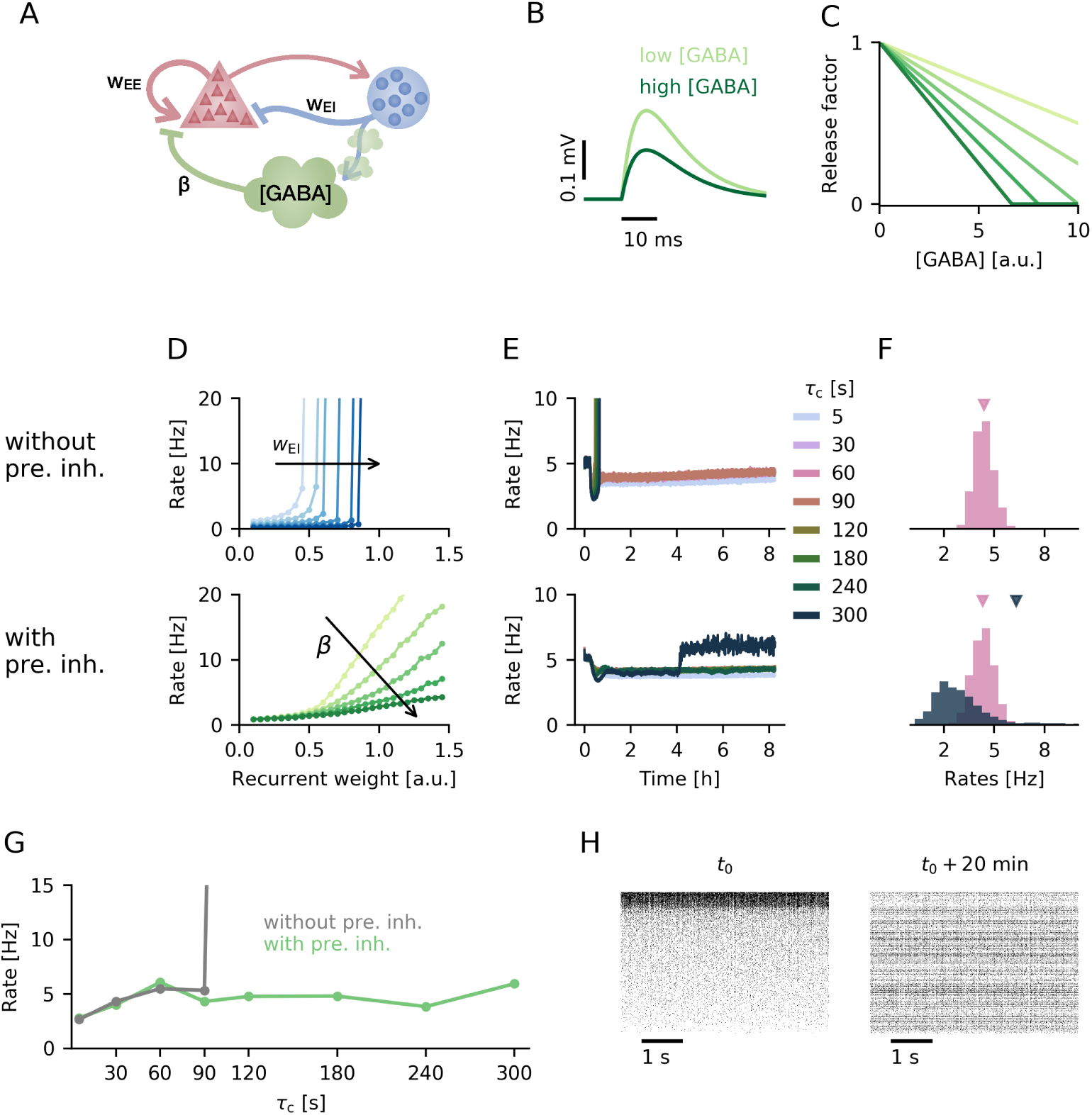
Presynaptic inhibition in a spiking network. **A.** Circuit diagram of the spiking network model consisting of an excitatory and inhibitory population of neurons as well as accumulation of local GABA concentration that triggers presynaptic inhibition. **B.** EPSP amplitude in the presence of presynaptic inhibition (*β* = 0.1) for low and high GABA concentration (1 and 5, respectively). **C.** Release factor as a function of GABA concentration for different examples of linear transfer functions. **D.** Population firing rate as a function of mean excitatory recurrent weight for increasing strengths of postsynaptic inhibition *w*_EI_ (top) or presynaptic inhibition *β* (bottom). **E.** Temporal evolution of the population firing rate for different homeostatic time constants *τ*_c_ in networks without (top) and with presynaptic inhibition (bottom). **F.** Final firing rate distribution for two example runs: *τ*_c_ = 60s (pink) and *τ*_c_ = 120s (dark blue-grey) without (top) and with presynaptic inhibition (bottom). Average firing rate is indicated by triangle of respective colour. **G.** Mean final population rate as a function of homeostatic time constant. **H.** Raster plot of 1000 random neurons in the networks sorted by activity at time *t*_0_ (left) and with the same ordering 20 minutes of simulation time later.

First we investigate how the mean firing rate depends on the overall excitatory recurrence. As in the rate model, networks without presynaptic inhibition exhibit a high sensitivity to the strength of recurrent excitatory weights (Fig. 4D, top) [5, 11]. For weak recurrent excitation the network fires at low rates, whereas for stronger excitation it destabilises and fires at a pathologically high firing rate that is only limited by the refractory period. Stronger postsynaptic inhibition merely shifts the point of destabilisation to higher values of excitatory recurrence. In contrast, networks with presynaptic inhibition allow a gradual increase of the mean population firing rate with growing excitatory recurrence (Fig. 4D, bottom). The dependence of the population firing rate on excitatory recurrent weight scales multiplicatively with the strength of presynaptic inhibition (Fig. 4C). The qualitatively different behaviour in response to changes in excitatory recurrence for networks with and without presynaptic inhibition is in good agreement with the results obtained in the rate network (cf. Fig. 1).

To verify whether presynaptic inhibition also stabilises spiking networks subject to Hebbian plasticity, we incorporate the triplet rule with homeostatically controlled, activity-dependent long-term depression [38] (see Methods and Models). The timescale of homeostasis is related to the time constant of the neuron-specific rate detector that scales the amount of long-term depression [60]. As previous work by Zenke et al. [60] has shown, stability in spiking networks without presynaptic inhibition critically depends on this homeostatic timescale. While fast homeostasis keeps the population firing rate close to target, slow homeostasis above a critical threshold is unable to control the positive feedback loop of recurrent excitation and Hebbian plasticity, such that the network exhibits runaway activity (Fig. 4E and F, top). Note that we observe higher critical homeostatic time constant than Zenke at al. [60]. This is a consequence of a higher background input in our spiking network (cf. Fig. 3) and may also depend on other differences in model choice, such as current-based rather than conductance-based synapses. Including presynaptic inhibition in the spiking network produces qualitatively similar results as in the rate model: Presynaptic inhibition increases the critical homeostatic time constant below which network activity is homogeneous and at the target rate (Fig. 4E, bottom). It also maintains low population firing rates beyond this critical timescale (Fig. 4G), although the network shows a broader distribution of single neuron firing rates (Fig. 4F). For slow homeostatic feedback control, we observe a similar turnover as in the rate network: the set of neurons that are most active changes over the duration of the simulation (Fig. 4H).

We conclude that the results we obtained for rate models can be qualitatively reproduced in networks of spiking neurons that include presynaptic inhibition based on local GABA spillover. In consequence, the features of presynaptic inhibition that prevent runaway excitation in the presence of Hebbian plasticity generalise across network models on different levels of abstraction.

## Discussion

In the present work we have demonstrated that presynaptic inhibition can robustly alleviate destabilisation of network activity caused by changes in recurrent excitation. In our model of presynaptic inhibition, synaptic transmission at excitatory synapses is inhibited presynaptically by GABA spillover from inhibitory synapses. Increases in recurrent excitation inevitably elevate inhibitory population activity and thus GABA release, forming a negative feedback loop on effective recurrent excitatory weights. This negative feedback loop serves as a rapid compensatory process that attenuates synaptic transmission if network activity is too high. Therefore, presynaptic inhibition prevents runaway excitation arising from Hebbian plasticity when homeostatic control mechanisms alone are not fast enough. By limiting the overall population activity, presynaptic inhibition allows for homeostasis on biologically realistic timescales without compromising network stability.

### Multiplicative gain control

We model presynaptic inhibition of synaptic transmission as a multiplicative factor onto excitatory recurrent synapses, which decreases with (inhibitory) network activity. Thus, presynaptic inhibition can serve as a form of multiplicative inhibition, i.e., a gain control. This was previously acknowledged by Fink et al. [18], who demonstrated in a simple feedback model that presynaptic inhibition at the neuro-muscular junction can control sensory gain. Yet, multiplicative inhibition of neural responses has typically been attributed to different processes such as shunting inhibition [31] or short-term depression [3].

Whether shunting inhibition – an increase in membrane conductance that short-circuits excitatory currents – can provide multiplicative inhibition [7] has been debated: Experimental and theoretical studies have shown that the overall effect of shunting inhibition on neural responses in fact is not multiplicative but subtractive [1, 12, 21], unless specific assumptions such as dendritic saturation or certain noise levels are met [12, 39]. Similar to the presynaptic inhibition mechanism studied here, short-term plasticity alters synaptic transmission by affecting transmitter release [28, 50]. Indeed is has been shown, that short-term depression can provide dynamic gain control [3, 40]. A defining difference between presynaptic inhibition and short-term plasticity is that presynaptic inhibition depends on surrounding network activity, whereas short-term plasticity is driven by presynaptic activity in the synapse in question. Hence, while presynaptic inhibition scales recurrent interactions globally and preserves the overall network structure, short-term depression reduces the impact of highly active neurons on the network.

We conclude that although shunting inhibition as well as short-term depression have been linked to multiplicative gain control, their effects are fundamentally different from presynaptic inhibition.

### Role of presynaptic inhibition in learning

Here, we have focused on the potential of presynaptic inhibition to stabilise Hebbian plasticity at recurrent excitatory synapses. It conserves the underlying network connectivity and might therefore set the stage for stable learning. GABA_B_ receptors are highly expressed in several regions traditionally linked to learning and memory [8, 41] and presynaptic inhibition is prominent in the hippocampus (see for example [25, 44, 48, 58]). It is therefore conceivable that presynaptic inhibition plays other roles in learning and memory beyond the stabilisation of ongoing plasticity demonstrated here. For example, experiments in hippocampus suggest that local, rapid and activity-dependent regulation of release probability serves to maintain synapses in an operational range, ensuring that synapses are optimally placed to undergo changes induced by learning mechanisms [10]. In addition, many models of learning rely on a coincidence of pre- and postsynaptic activity (i.e., Hebbian plasticity). By regulating information flow, presynaptic inhibition could thus act as a gating mechanism for the induction of plasticity [34, 53]. Indeed, defects in GABA_B_ receptor expression have been shown to compromise long-term plasticity, leading to impairments in hippocampus-dependent memory [54].

Gating of long-term plasticity by presynaptic inhibition was also observed in the amygdala, where loss of presynaptic GABA_B_ receptors led to a generalisation of conditioned fear to non-conditioned stimuli [42]. In this context it was suggested that presynaptic inhibition sets the balance between associative and non-associative long-term potentiation. In cerebellum, stimulation with physiological activity patterns leads to changes in presynaptic GABA_B_ receptor expression, which was suggested to complement other forms of plasticity: A reduction in presynaptic inhibition increases synaptic transmission and could thus enhance long-term plasticity [36].

In summary, a range of experiments indicate that beyond providing stability during ongoing plasticity, presynaptic inhibition could serve as an activity-dependent gating mechanism for long-term plasticity.

### Phenomenology and limitations

Because our goal was to investigate consequences of presynaptic inhibition on the network level, we adopted a phenomenological description of the mechanism. Although we motivated the model by a specific pathway that suppresses presynaptic calcium channels, activation of presynaptic GABA_B_ receptors can impair the release machinery in other ways, e.g. by activation of potassium channels [58]. Furthermore, presynaptic terminals also express GABA_A_ receptors that have been implicated in presynaptic inhibitory effects [17, 57]. Such alternative pathways also operate on timescales in a sub-second range, and should therefore not influence the validity of the mechanisms we suggest. Furthermore we show that the compensation of recurrent excitation does not require presynaptic inhibition to exclusively target recurrent excitatory connections (SI Appendix, Fig. S1), supporting the generalisability of the mechanism.

Urban-Ciecko et al. have attributed presynaptic inhibition to somatostatin-positive (SOMs) but not parvalbumin-positive interneurons (PVs) [53]. Modelling circuits with different interneuron types, in which presynaptic inhibition is mediated by SOMs is beyond the scope of this work, however.

For analytical tractability we mostly used linear functions for presynaptic inhibition, but show that the stabilising effect is robust to the specific choice of transfer function (SI Appendix, Fig. S4). Another simplifying assumption in our work is that we consider synaptic transmission to be deterministic, such that presynaptic inhibition merely affects a synaptic release factor. A natural extension would be to include a probabilistic release mechanism and a dynamic model of short-term plasticity. However, we expect that our results will still hold qualitatively, because on the slow timescales of synaptic plasticity, short-term plasticity will act mainly through changes in steady state [49], rather than through a short-term redistribution of synaptic release [29].

### Spatial specificity of presynaptic inhibition

It is unclear how specific is the mechanism of presynaptic inhibition. A critical determinant of this specificity is the source of GABA at excitatory synapses. One possibility is the presence of axo-axonal synapses that release GABA in very close proximity to excitatory synapses and thereby mediate a potentially highly specific form of presynaptic inhibition [27]. For example, presynaptic inhibition at the neuro-muscular junction ensures smooth and stable movement patterns through axo-axonal synapses [18]. However, in most brain areas axo-axonal synapses are not numerous enough to account for the full range of presynaptic inhibition effects [53]. Presynaptic inhibition also occurs in the absence of axo-axonal synapses, potentially through GABA that diffuses from nearby inhibitory synapses [16, 33]. In fact, a specific class of interneurons – neurogliaform cells – has been found to release GABA in a target-independent way, generating non-specific forms of inhibitory control [37]. Regardless of the exact source of GABA spillover, the spatial specificity of presynaptic inhibition depends on diffusion coefficient of GABA as well as the input specificity of the respective cells.

We were interested in the interactions of presynaptic inhibition with plasticity in recurrent connections. We therefore considered a small network in which synaptic connections can essentially be considered random. Assuming that GABA release and spillover provide a spatially unspecific signal on this scale, we modelled presynaptic inhibition as a global effect. More specifically, activity of all inhibitory neurons modulates a single release factor that acts at every recurrent excitatory synapse. This leads to the control of mean population activity by presynaptic inhibition, whereas single neurons can exhibit heterogeneous firing rates. However, we do not expect that in the brain, presynaptic inhibition can act sufficiently locally to allow highly specific control, e.g., on a single neuron level. The reason is that the mechanism does not provide direct feedback but acts through a population of inhibitory neurons, thus losing spatial specificity.

Considering morphologically more complex neurons might give insights into the computational properties of presynaptic inhibition at the level of single compartments or even dendritic branches. Experiments have revealed that release probabilities within a dendritic branch are similar and change depending on recent dendritic activity [10]. However, an extension to networks of multi-compartment neurons is also beyond the scope of this project.

In conclusion, the spatial specificity of presynaptic inhibition is not yet resolved. Both local, specific mechanisms via axo-axonal synapses and broader mechanisms mediated by GABA spillover from other synapses have been demonstrated [4]. It is also conceivable that the degree of specificity of presynaptic inhibition varies between brain areas depending on the computational demands.

### Outlook

In the present work, we focused on the role of presynaptic inhibition to stabilise recurrent excitation in the presence of plasticity. Besides the extensions mentioned in previous paragraphs, it would also be interesting to study the effects of presynaptic inhibition on sensory information processing, e.g., by providing a gating mechanism for sensory information at an early stage [4]. Presynaptic inhibition has been repeatedly observed in early sensory systems including the retina [47], lateral geniculate nucleus [13], somatosensory cortex [53] and the olfactory system of drosophila [35], and theoretical work has implicated presynaptic inhibition in extending the dynamic range in sensory processing [61]. Further theoretical work will be required to pinpoint the full functional repertoire of presynaptic inhibition.

## Methods and Models

To investigate the effect of presynaptic inhibition on network stability, we simulate rate-based and spiking recurrent networks and mathematically analyse mean population dynamics.

### Recurrent rate network model

The recurrent rate network consists of 1024 excitatory and 256 inhibitory neurons, which are randomly connected and described by their firing rates. The firing rate dynamics are given by

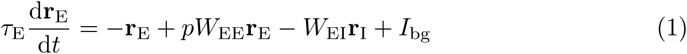

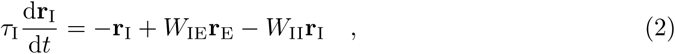

where **r**_E_ and **r**_I_ are vectors of excitatory and inhibitory neuron rates, *W*_*xy*_ with *x, y ∈* {E, I} are matrices of synaptic weights and *I*_bg_ is the background input to excitatory neurons. In addition, firing rates are rectified to ensure positivity. The timescale of rate dynamics is dictated by *τ*_E_ and *τ*_I_ (see SI Appendix, Methods and Models for parameter values).

Presynaptic inhibition is implemented as a global release factor *p* that multiplicatively scales the total excitatory input and which is modulated by the total inhibitory firing rate 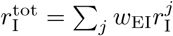 according to

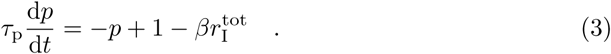

The release probability is also rectified. The parameter *β* determines the decrease in release factor with inhibitory firing rate and thus regulates the strength of presynaptic inhibition.

We used the Bienenstock-Cooper-Munro (BCM) [6] rule as a model for synaptic plasticity of the recurrent connections of excitatory neurons, with an effective timescale *τ* ^plast^ = 60s. This Hebbian rule contains a sliding threshold that determines the direction of weight changes based on a delayed running estimate of single neuron firing rates. If the threshold is updated on sufficiently short timescales, it can act as a homeostatic mechanism that brings single neurons to a target rate (here 5 Hz) [60]. The time constant of this threshold is denoted as homeostatic (or control) timescale *τ*_c_ and critically determines network stability.

### Critical homeostatic timescale

In order to understand how presynaptic inhibition impacts the range of homeostatic timescales for which the networks shows stable firing at the target rate, we analyse the stability of a mean population model derived from the full network. Using a separation of timescales, we reduce the system to two dimensions, conduct a linear stability analysis and finally obtain an analytical expression for the critical homeostatic timescale – the maximal timescale of the homeostatic component of the BCM rule for which we expect the network to remain stable:

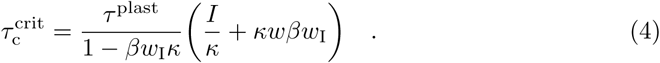

This critical timescale depends on presynaptic inhibition strength *β*, background input *I*, target rate *κ*, overall inhibitory feedback *w*_I_ and the level of recurrence *w* (see SI Appendix, Methods and Models for the derivation). Note that the absolute value of the timescale needs to be seen relative to the timescale of plasticity *τ* ^plast^.

### Spiking network model

The recurrent spiking network consists of 4000 excitatory and 1000 inhibitory leaky integrate-and-fire neurons, which are randomly connected with current-based synapses. Excitatory neurons receive additional background input from Poisson units. As in previous work [60], synaptic plasticity and homeostasis are modelled by the triplet rule [38] – the spiking version of the BCM rule [19] – with delayed rate estimation (see SI Appendix, Methods and Models for details and parameters).

## Supporting Information

### Methods and Models

To investigate the effect of presynaptic inhibition on network stability, we simulate large rate-based and spiking recurrent networks and mathematically analyse mean population dynamics.

#### Recurrent rate network

The recurrent rate network consists of *N*_E_ = 1024 excitatory and *N*_I_ = 256 inhibitory neurons described by their firing rates 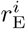 and 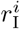. The neurons are randomly connected with connection probability *c* = 100 × 2^−10^ ≈ 0.1 and fixed indegree. Neuron numbers were chosen as powers of two to increase simulation speed, and the connection probability guarantees an exact ratio of 4 : 1 (100 and 25) excitatory and inhibitory ingoing synapses at each neuron. The firing rate dynamics are given by

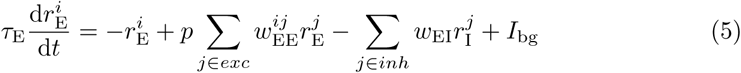

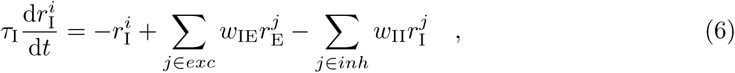

with synaptic weights 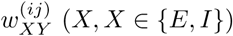 and time constants *τ*_E_ = *τ*_I_ = 20 ms. Presynaptic excitatory or inhibitory input populations are given by indices from neuron-specific subsets *exc* and *inh* and excitatory neurons receive noisy, uncorrelated background input *I*_bg_ (with distribution *𝒩* (*I*, 0.1*I*)). To ensure positive and finite firing rates, 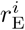 and 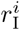 are rectified and saturated at 200 Hz. Synaptic weights are scaled by connection probability and network size (see Table 1) and fixed with exception of the excitatory recurrent weights 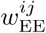. Excitatory weights are either a free parameter (Fig 1) or subject to plasticity. Weights are chosen such that the network is balanced [11] and excitatory and inhibitory firing rates are similar.

**Table 1.** Rate network parameters. (unless specified differently)

| parameter | value | parameter | value |
| --- | --- | --- | --- |
| $N_{\text{E}}$ | 1024 | $w_{\text{EI}}$ | $1/(cN_{\text{I}})$ |
| $N_{\text{I}}$ | 256 | $w_{\text{IE}}$ | $1.5/(cN_{\text{E}})$ |
| $c$ | $100 \times 2^{-10}$ | $w_{\text{II}}$ | $0.5/(cN_{\text{I}})$ |
| $r_{\text{max}}$ | 200 Hz | $I$ | 0.5 Hz |

#### Presynaptic inhibition mechanism

The global release factor *p* in Eq (5) multiplicatively scales the total excitatory recurrent input to each neuron. In the absence of presynaptic inhibition we set *p* ≡ 1, which leaves a standard recurrent rate model of excitatory and inhibitory neurons. If the presynaptic inhibition mechanism is included, *p* is modulated by the total inhibitory input rate according to

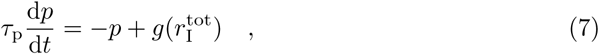

where *g* is a monotonically decreasing function and *τ*_p_ = 500 ms. For simplicity, we use a linear rectified function given by *g*(*r*) = [1 − *βr*]_+_ with slope *β*. The mechanism is global, meaning that a single release factor *p* is modulated by the total weighted inhibitory activity 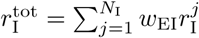. The slope *β* as well as inhibitory weight *w*_EI_ determine the strength of the mechanism. To account for the scaling of *w*_EI_, *β* is also normalised by connection probability *c* (see Table 2).

**Table 2.** Timescales, presynaptic inhibition and plasticity parameters in the rate network. (unless specified differently)

| timescale | value | parameter | value |
| --- | --- | --- | --- |
| $\tau_E = \tau_I$ | 0.02 s | $\eta$ | 5 |
| $\tau_P$ | 0.5 s | $\kappa$ | 5 Hz |
| $\tau_P$ | 300 s | $w_{\text{max}}$ | 10 $w_0$ |
| $\tau_c$ | free parameter | $\beta$ | 0.05 c (default) |

In **??** we test sigmoid and exponential functions as alternatives. The sigmoid transfer function is parametrised as

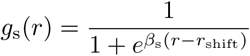

with slope parameter *β*_s_ and shift (or reversal point) *r*_shift_. For an exponentially decaying transfer function we use

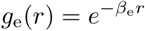

with slope parameter *β*_e_.

#### Plasticity model

We used the Bienenstock-Cooper-Munro (BCM) rule [6] as a model for synaptic plasticity. Plasticity only affects connections between excitatory neurons, changing the weights according to

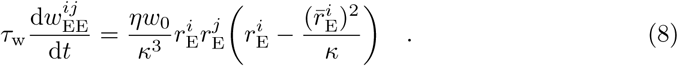

The effective timescale of plasticity is determined by time constant *τ*_w_, initial weight *w*_0_ and learning rate *η*. Thus, the effective time constant of plasticity is given by

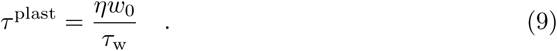

As in the original formulation, this recurrent BCM rule is Hebbian and will attempt to potentiate afferent synapses of neurons above the target rate *κ* and depress the ones below, which is captured by the nonlinear threshold 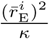 in Eq (8). The neuron-specific threshold depends on the target rate *κ* as well as a running estimate 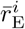 of neuron activity, given by a low-pass filter of that neuron’s firing rate:

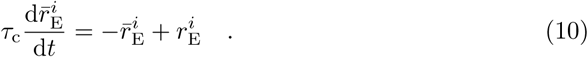

For constant input, the plasticity rule together with the threshold’s dependence on the rate estimate create a homeostatic system that tries to bring the firing rate of every neuron to the target *κ* = 5 Hz. The network is initialised close to the fixed point as determined by mathematical analyses: 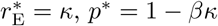 and 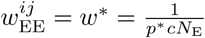 (cf. Eq (22)). Excitatory recurrent weights are limited to the interval from zero to 10*w\**.

A critical parameter for stability is the time constant *τ*_c_ of rate estimation [60], which in this context acts as timescale of homeostasis and is a free parameter of the system. Note that the homeostatic timescale needs to be seen in relation to the the effective timescale of plasticity. To improve simulation speed in the rate network, we rescaled the timescale of plasticity by a factor of 50 (*τ* ^plast^ = 60s instead of 3000s used in previous work [60]). As all other timescales in the system are at least a factor of 100 faster, this does not influence the stability of the network. Thus, a homeostatic timescale of 60 seconds (*τ* ^plast^) in our network actually corresponds to homeostasis on the order of almost an hour of biological time.

#### Mean population rate model

To understand the dynamics of the recurrent rate network in more detail, we derive a mean population rate model from the full network. Assuming homogeneous neuron activity as well as a sufficiently large network, the excitatory and inhibitory populations can be described by their mean population firing rate *r*_E_ and *r*_I_, respectively. Both populations receive recurrent and reciprocal input with a mean weight, such that their dynamics can be simplified to

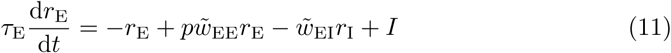

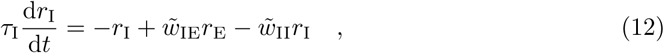

with weights 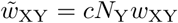 and in particular 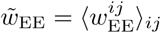, where *w*_XY_ are the synaptic weight parameters from the full rate network (Table 1). As in the full model, firing rates are rectified and thresholded at 200 Hz and presynaptic inhibition is modelled by a monotonic decrease in release factor *p* (see Eq (7)) with 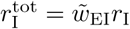. As 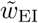 does not depend on the connectivity or number of neurons, no further normalisation of *β* is needed.

In the mean population rate model the BCM rule with homeostatically sliding threshold reduces to

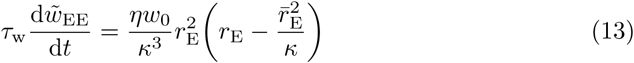

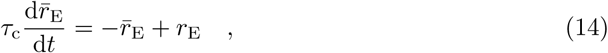

as pre- and postsynaptic firing rates are both described by the mean excitatory population rate. Thus, the mean population rate model with presynaptic inhibition simplifies to a five-dimensional dynamical system.

#### Analytical steady-state firing rates

To compare the steady-state behaviour of the mean population rate model in the presence and absence of presynaptic inhibition, we derive the fixed point of Eq (11) and (12) for unsaturated rates and linear presynaptic inhibition transfer function in Eq (7). In steady-state, the inhibitory rate is

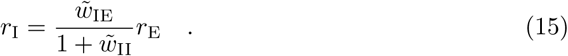

and thus we can write the steady-state excitatory firing rate without presynaptic inhibition as

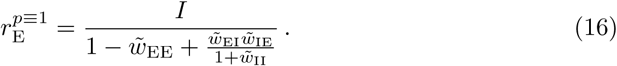

Note that this fixed point only exists as long as 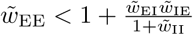.

In the following we define the total inhibition recruited by the excitatory population through the inhibitory population, and the excitatory weight as

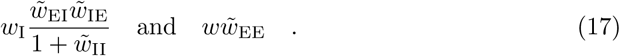

First, we assume that the mean background input *I* is weak enough to ensure that *r*_E_ *≤* 1*/*(*w*_I_*β*). In this regime the transfer function *g* is linear such that Eq (11) can be rewritten to

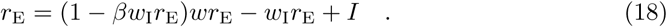

Solving this quadratic equation for *r*_E_ gives the steady-state firing rate with presynaptic inhibition for linear transfer function:

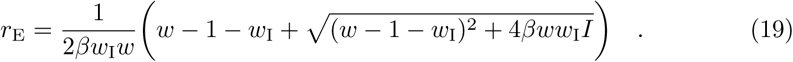

Note that the solution with a negative contribution of the square root does not give positive firing rates and thus is not relevant for this system.

It can be shown that for mean background input 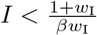, presynaptic inhibition imposes an upper bound on the steady-state firing rate with respect to the recurrent excitatory weight *w*. Taking the limit of *w* to infinity gives

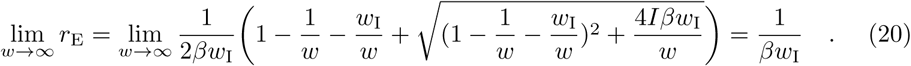

Therefore, the upper bound of the excitatory firing rate for moderate external inputs depends on the strength of presynaptic inhibition *β* as well as the total effective inhibitory weight *w*_I_ (see definition in Eq (15).

For external input strengths 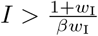, the excitatory rate *r*_E_ is larger than 1*/*(*βw*_I_) leading to *p* = [1 − *βw*_I_*r*_E_]_+_ = 0. In this case the steady-state firing rate reduces to

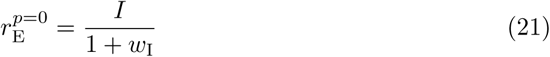

and hence is independent of the excitatory recurrent weight *w*.

#### Derivation of critical homeostatic timescale

To derive the critical homeostatic timescales in the presence and absence of presynaptic inhibition, we revisit the five-dimensional dynamical system given by Eq (7) and (11)-(14), and follow a similar approach as Zenke et al. [60]: first we reduce the system to two dimensions, then perform a linear stability analysis around the fixed point and finally determine how the stability of the fixed point depends on the timescale of homeostasis.

#### Reduction to two-dimensional system

As the dynamics of *r*_E_, *r*_I_ and *p* are much faster than those of *w* and 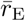 (i.e., *τ*_E_, *τ*_I_, *τ*_*p*_ ≪ *τ*_w_, *τ*_c_), we can use a separation of timescales approach to reduce the full system to two dimensions. With respect to the slow plasticity and homeostasis dynamics, the fast variables are at their steady-state. For the excitatory firing rate we can therefore write

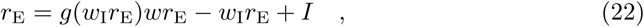

where we again use the definitions of *w* and *w*_I_ from Eq (17) for readability. Taking the derivative with respect to *w* on both sides gives

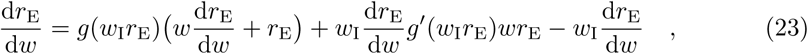

where *g′* is the derivative of *g* with respect to *r*_E_. Solving for 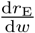 leads to

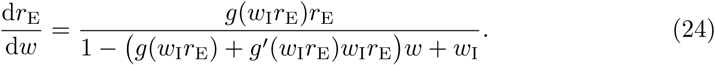

To eliminate *w* from the partial derivative above we reorder Eq (22) for *w*, insert the result and after simplifying the expression finally obtain

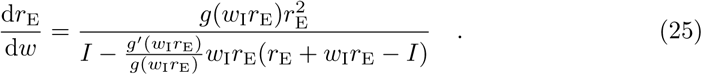

Using the chain rule 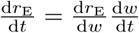, we can express slow dynamics of the BCM-type plasticity rule in Eq (13) through its effect on the steady-state of *r*_E_. Thus the firing rate *r*_E_ becomes the slow dynamic variable together with its running estimate 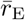 and therefore the final two-dimensional system is described by

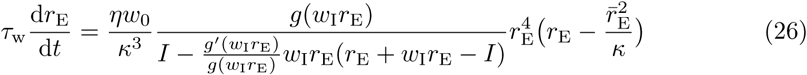

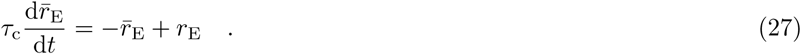

#### Linear stability analysis

The system described by Eq (26) and (27) is in the stationary state if the rate estimate and excitatory firing rate are at the target 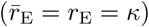. To obtain the Jacobian of the system, we calculate the partial derivates of the right hand side of Eq (26) and Eq (27) with respect to *r*_E_ and 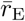 and evaluate it at the fixed point 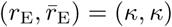:

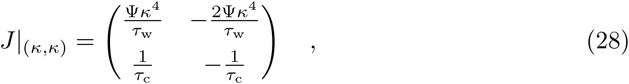

where we introduced the auxiliary expression

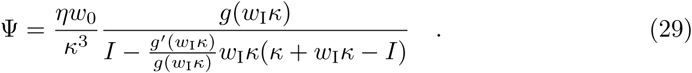

The characteristic polynomial then is

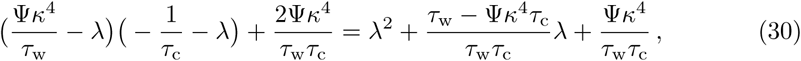

which determines the eigenvalues of the system linearised around the fixed point:

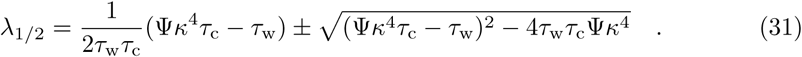

If the real part of both eigenvalues is negative, the fixed point is stable (see for example [23]). In the following we use analogous reasoning to previous work [60] to prove that this condition is fulfilled as long as 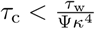. First of all, Ψ is positive for moderate external input (*I* < *κ*(1 + *w*_I_)), because *g* is a monotonously decreasing function and all other variables are positive. The stability condition follows immediately if the square root is imaginary. In case the square root is real, we can write the larger eigenvalue as

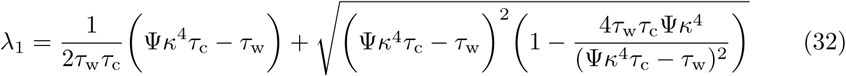

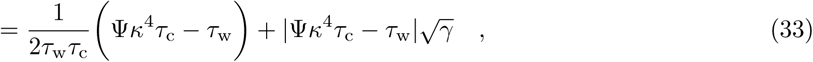

where *γ* is the second term of the product in the square root and 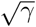 has to be real given the previous assumption. If Ψ*κ*^4^*τ*_c_ > *τ*_w_ then 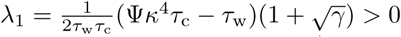 and thus the fixed point is unstable.

If, on the other hand, Ψ*κ*^4^*τ*_c_ < *τ*_w_ it follows that

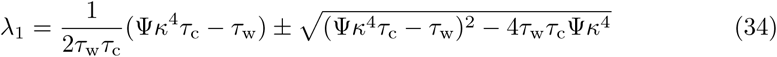

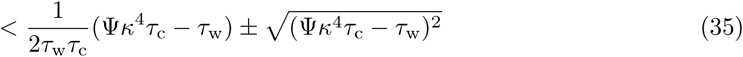

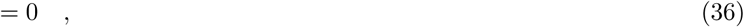

such that both eigenvalues are negative and therefore the fixed point is stable.

#### Critical homeostatic timescale

Using the stability condition derived above, we can identify the critical homeostatic timescale as

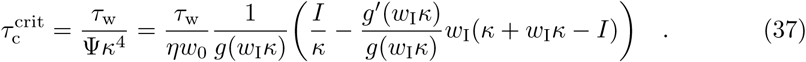

To obtain a more intuitive expression, we reorder Eq (22) at the fixed point (*r*_E_ = *κ*) to give *g*(*w*_I_*κ*)*wκ* = *κ* + *w*_I_*κ* + *I* and insert this into the Eq (37) to get

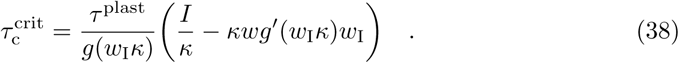

In summary, we have shown that stability of the fixed point at the target rate *κ* is guaranteed as long as the homeostatic time constant *τ*_c_ is smaller than a critical value 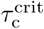, which depends on the release factor at the target rate *g*(*w*_I_*κ*), excitatory recurrence *w*, the change *g′*(*w*_I_*κ*) of release factor with inhibitory acitvity (i.e. GABA spillover) and the dependence *w*_I_ of this inibitory activity on excitatory firing rate. Note that the critical homeostatic timescale needs to be seen in relation to the effective time constant of plasticity 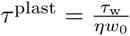.

Without presynaptic inhibition the two-dimensional system simplifies to

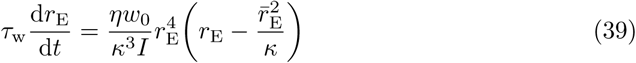

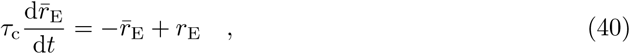

which was previously analysed by Zenke et al. [60]. We can recover their results by inserting *g*(*r*) ≡ 1 (implying *g′*(*r*) ≡ 0) into Eq (37), which leads to the critical timescale without presynaptic inhibition

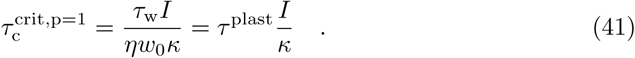

**??** shows the eigenvalues for increasing homeostatic time constant as well as the phase plane dynamics with fast and slow homeostasis, both with and without presynaptic inhibition.

#### Spiking network model

The recurrent spiking network we studied consists of 5000 leaky integrate-and-fire neurons (4000 excitatory and 1000 inhibitory neurons), which are randomly connected with current-based synapses [55]. In addition to the recurrent connections, both excitatory and inhibitory neurons receive random background input from an external population modelled by 2000 Poisson units with firing rate 5 Hz. Connection probability for recurrent and external connections is sparse (10%).

The membrane potential *V*_*i*_ of neuron *i* evolves according to

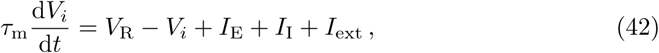

with reversal potential *V*_R_, membrane time constant *τ*_m_ and synaptic input currents from excitatory (*I*_E_), inhibitory (*I*_I_) and external populations (*I*_ext_). If a voltage trace crosses the threshold *V*_T_, the neuron emits a spike and the voltage is reset to *V*_R_ for an absolute refractory period of *τ*_ref_ (see Table 3 for an overview of neuron and network parameters). The spike train of a single neuron *i* is defined as a sum over spikes *k* given by 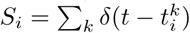 with spike times 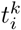. Input currents are given by a temporally filtered, linear, weighted sum of the spikes from the respective population to neuron *i*:

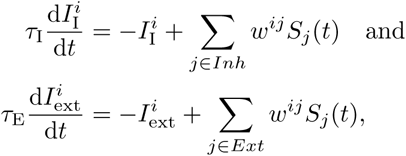

where *w*^*ij*^ are synaptic weights from neuron *j* to neuron *i*.

**Table 3.** Neuron and network parameters.

| neuron parameter | value | network parameter | value |
| --- | --- | --- | --- |
| $V_R$ | -70 mV | $N_E$ | 4000 |
| $V_T$ | -50 mV | $N_I$ | 1000 |
| $\tau_m$ | 20 ms | $N_{\text{ext}}, r_{\text{ext}}$ | 2000, 5 Hz |
| $\tau_{\text{ref}}$ | 5 ms | connectivity $c$ | 0.1 |

#### Presynaptic inhibition by GABA spillover

In analogy to the rate network, the effect of presynaptic inhibition on synaptic transmission is modelled as a multiplicative factor (“release factor”) onto excitatory synaptic weights. However, in the spiking network we consider a more local version of the mechanism with neuron-specific release factors *p*^*i*^ that are modulated by GABA spillover. More specifically, excitatory synaptic inputs follow

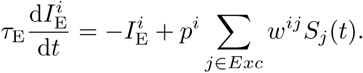

We estimate local GABA levels (can also be interpreted as concentration) by accumulating a fixed amount of GABA - given by *A*_GABA_ - for every inhibitory spike according to

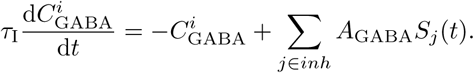

The local GABA levels are neuron-specific and affected by all inhibitory neurons projecting to the respective neuron. Thus, an inhibitory spike leads to both an inhibitory current and an increase in local GABA level at downstream excitatory neurons. The inhibition of release through GABA spillover is then again modelled by a linear decrease in release factor with increasing GABA levels, that is

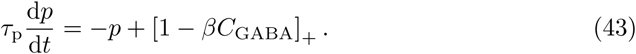

We set the time constant of this process to *τ*_p_ = 300 ms to roughly match the timescale of presynaptic inhibition observed experimentally [53]. As before, the slope *β* determines the strength of presynaptic inhibition and *p* ≡ 1 recovers the traditionally used current-based network.

Although we consider *C*_GABA_ to be related to local GABA concentration, it is unitless and can not be mapped to any specific biophysical quantity. Nevertheless, we normalise *A*_GABA_ by the number of ingoing inhibitory synapses, such that the local GABA level *C*_GABA_ roughly relates to the inhibitory firing rate. This allows us to use a presynaptic inhibition strength paremeter *β* on the same order of magnitude as in the rate and mean population model (without further scaling). Table 4 contains a parameter summary for presynaptic inhibition and the synaptic weights. We omitted the units of the weights as they have no biophysical meaning in this simple current-based spiking model. Note that for consistency in the units of Eq(42) the unit would theoretically have to be Volt.

**Table 4.** Synaptic connection and spillover parameters.

| connection | value | parameter | value |
| --- | --- | --- | --- |
| $w_{EE}$ | free parameter | $w_{\text{ext}}$ | 2.0 |
| $w_{IE}$ | 0.5 | $A_{\text{GABA}}$ | $1/(cN_I)$ |
| $w_{EI}$ | -2.75 | $\beta$ | 0.1 or free parameter |
| $w_{II}$ | -2.75 | $\tau_p$ | 300 ms |

#### The triplet plasticity rule

We model plasticity of excitatory synapses in the spiking network using the minimal triplet STDP model tuned to visual cortex data [38] and metaplasticity implemented by homoeostatically regulating the amount of long-term depression (LTD) depending on the postsynaptic firing rates [38, 60]. The amplitude of LTD is given by

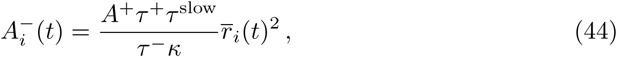

where *A*^+^ is the amount of long-term potentiation (LTP), *τ* ^+^, *τ* ^slow^ and *τ*^−^ timescales of the triplet STDP model (see [60]) and *κ* the target rate at which the definition of 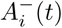 ensures balance of LTD and LTP. Changes in the LTD amplitude of neuron *i* are driven by the moving average of its postsynaptic firing rate

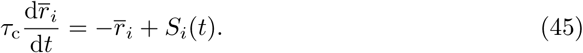

The overall timescale of this homoeostatic component of the metaplastic triplet STDP rule is therefore given by *τ*_c_ and as in the rate network a crucial parameter for network stability [60].

Weight modifications at excitatory recurrent synapses are limited to the interval 0 < *w*_ij_ < *w*_max_ during ongoing plasticity, whereas structural changes are not allowed (synapses initialised at zero strength remain absent). A summary of parameters related to plasticity is shown in Table 5. The effective time constant of plasticity can be approximated in a mean field model from the plasticity parameters by considering the expected mean weight update. Given the default parameters of this network, the effective timescale of plasticity amounts to *τ*_w_ = 2975 s (see [60] for details).

**Table 5.** Plasticity parameters.

| parameter | value | parameter | value |
| --- | --- | --- | --- |
| $A^+$ | $6.5 \times 10^{-3}$ | $\kappa$ | 5 Hz |
| $\tau^+$ | 11.8 ms | $w_0$ | 1(0.5 without pre.inh.) |
| $\tau^{\text{slow}}$ | 114 ms | $w_{\text{max}}$ | $5w_0$ |
| $\tau^-$ | 33.7 ms | $\tau_c$ | free parameter |

#### Numerical simulations

The numerical equations of the rate network were integrated using forward Euler with a timestep of 1 ms. Simulations were run in Python with support of Cython for speed improvement.

Spiking network were simulated using the Brian2 package [46] for Python. Numerical equations were integrated using Runge-Kutta method of 4th order for the neuron equations and forward Euler for synaptic interactions. The integration timestep was 0.1 ms.

**Fig S1.**
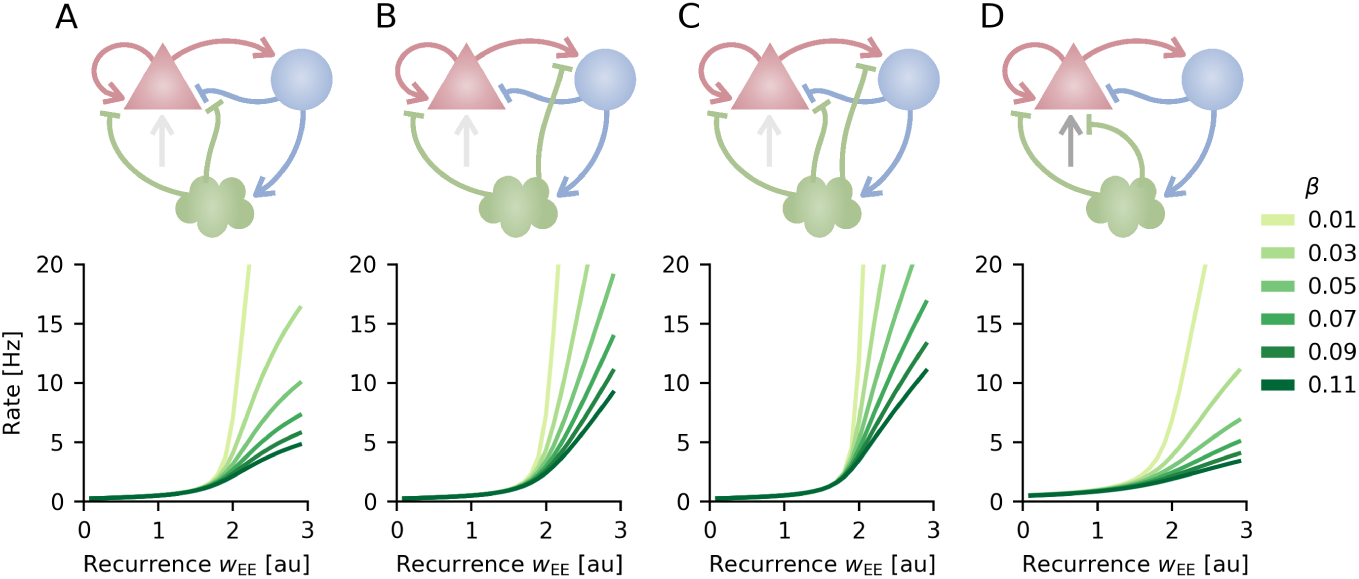
Presynaptic inhibition also compensates for increases in recurrent excitation in alternative circuit motifs. Top: Mean population models in which presynaptic inhibition not only affects excitatory recurrent synapses, but also **A.** inhibitory synapses onto excitatory cells, **B.** excitatory synapses onto inhibitory cells **C.**, all recurrent synapses (including recurrent inhibitory connections, not shown) or **D.** background input. Bottom: Firing rate of excitatory population in respective circuit model as a function of excitatory recurrence for increasing strengths of presynaptic inhibition (*β*).

**Fig S2.**
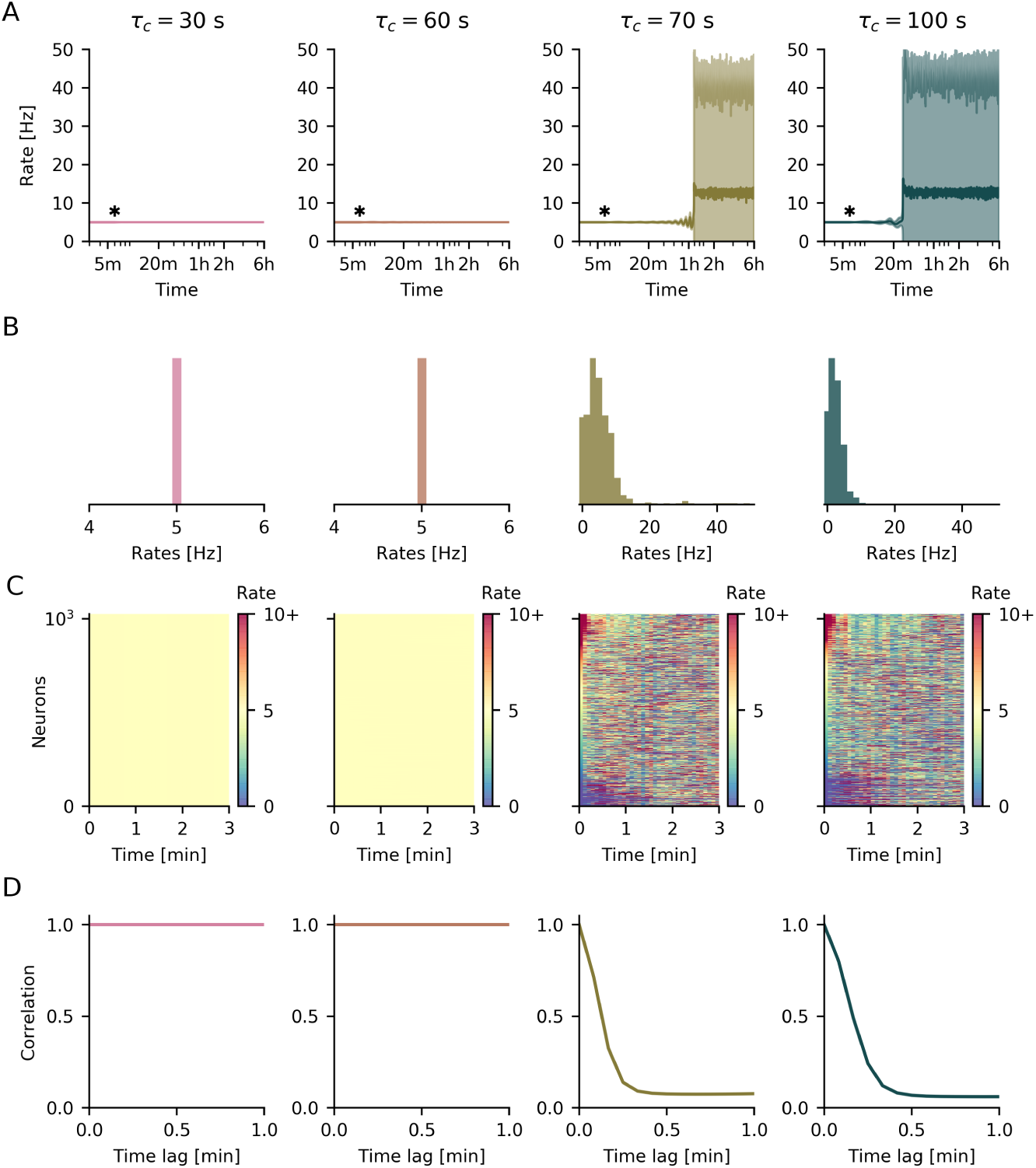
The homeostatic timescale affects the distribution and temporal correlation of single neuron firing rates. **A.** Mean (solid line) and standard deviation (shaded area) of firing rate over time. Plasticity is switched on after 6 minutes of simulation time (asterisk). **B.** Distribution of single neuron firing rates. **C.** Firing rate of single neurons over time. Target rate is indicated by yellow colour. **D.** Temporal correlation of single neuron firing rates.

**Fig S3.**
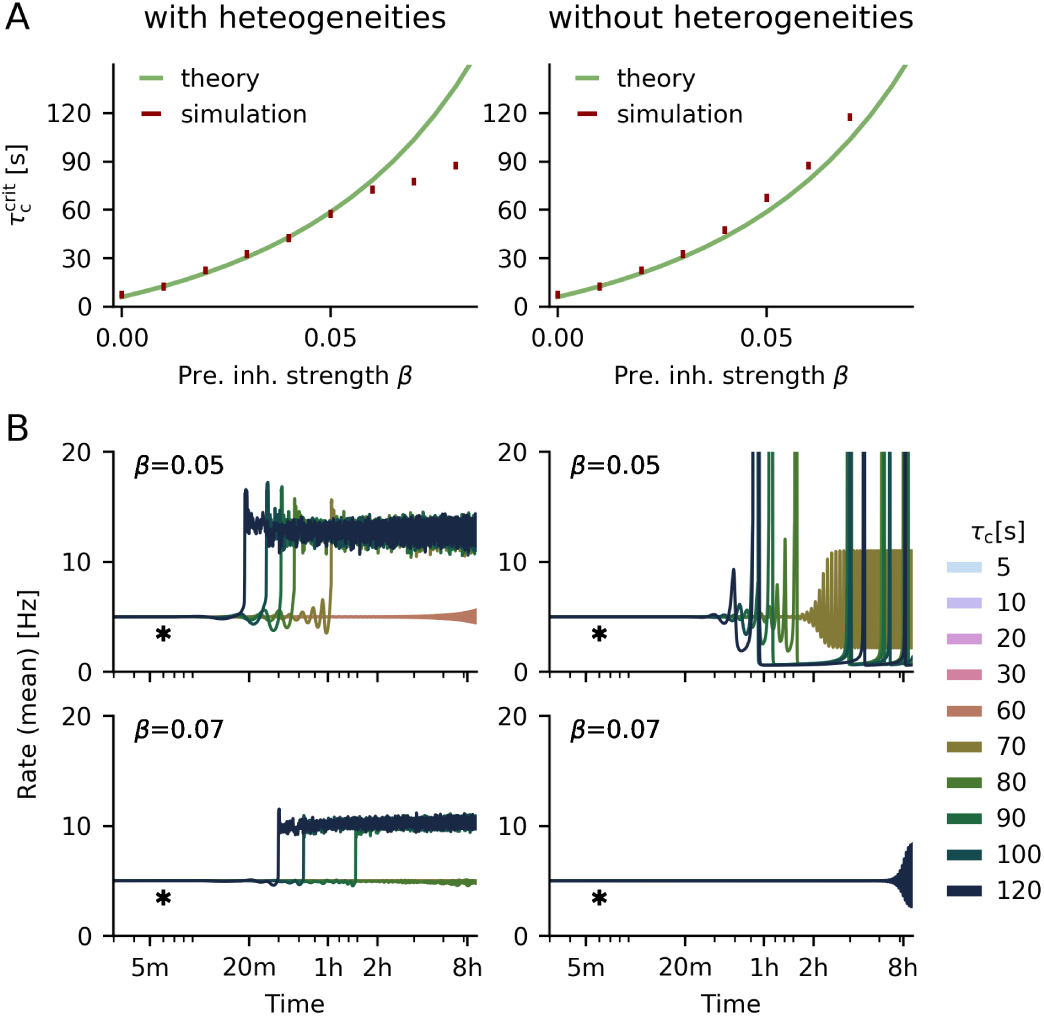
Removing heterogeneity (noise on initial excitatory recurrent weights) improves the fit of simulated and analytically predicted critical timescales. **A.** Critical homeostatic time scale as a function of the presynaptic inhibition strength *β* in networks with (left) and without (right) heterogeneity. For simulations the vertical bars indicate the range between largest stable and smallest unstable timescale tested. **B.** Temporal evolution of the average firing rate for different homeostatic time constants *τ*_c_ with (left) and without (right) heterogeneity in the network for different strengths of presynaptic inhibition *β*.

**Fig S4.**
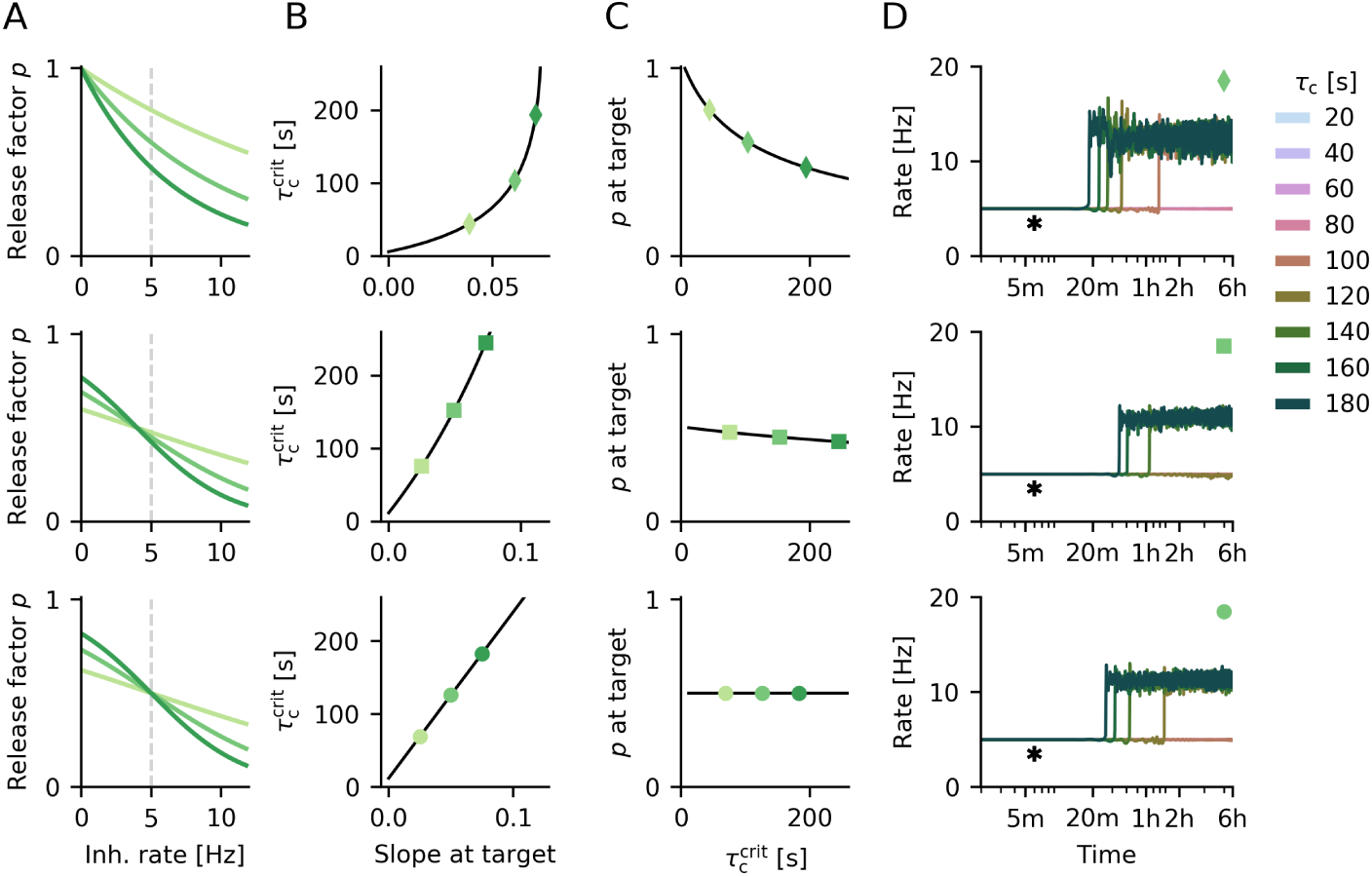
The stabilising effect of presynaptic inhibition is robust to the transfer function choice. **A.** Exponentially decreasing (top) and sigmoid transfer functions (bottom two) linking inhibitory firing rate to release factor for different slope parameters (*β*_e_ = 0.05, 0.1, 0.15, *β*_s_ = 0.1, 0.2, 0.3, shift of sigmoid: 4 (middle) and 5 (bottom). Target rate of 5 Hz is indicated by gray dashed line. **B.** Increase in analytical critical homeostatic timescale as a function of transfer function slope at the target rate. Parametrisation in A indicated by markers of same colour. **C.** Release factor at target rate as a function of increase in homeostatic time constant. Markers correspond to transfer functions shown in A. **D.** Temporal evolution of the average firing rate in networks with presynaptic inhibition for different homeostatic time constants *τ*_c_. Plasticity is switched on after 6 minutes of simulation time (asterisk). Coloured marker in top right corner specifies parametrisation of transfer function (see A).

**Fig S5.**
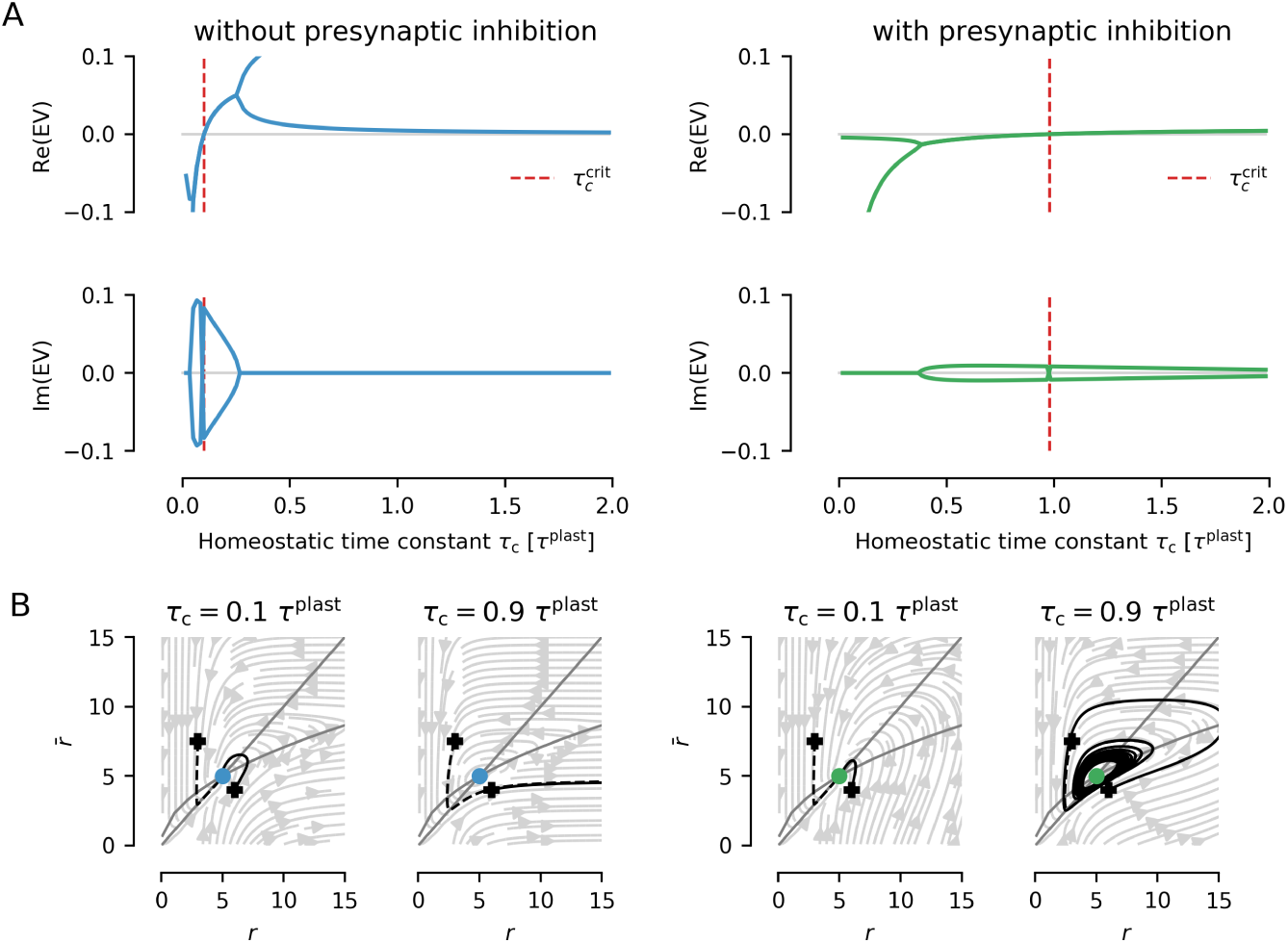
Linear stability in the reduced (two-dimensional) mean population model critically depends on the time constant of homeostasis. **A.** Eigenvalues of reduced system with compared to without presynaptic inhibition as a function of homeostatic time constant. The time constant is given in units of effective plasticity time constant *τ* ^plast^. The critical homeostatic timescale (where the eigenvalues become positive) is marked by a red dashed line. **B.** Phase plane dynamics with nullclines and example trajectories for short and long homeostatic timescales in systems with and without presynaptic inhibition.

## Acknowledgments

We thank Loreen Hertä g for helpful discussions and critical reading of the manuscript. We thank Owen Mackwood for considerable technical support with running simulations on a compute cluster.

## References

1. L. Abbott and F. S. Chance. Drivers and modulators from push-pull and balanced synaptic input. Progress in brain research, 149:147–155, 2005.

2. L. F. Abbott and S. B. Nelson. Synaptic plasticity: taming the beast. Nature neuroscience, 3(11):1178–1183, 2000.

3. L. F. Abbott, J. Varela, K. Sen, and S. Nelson. Synaptic depression and cortical gain control. Science, 275(5297):221–224, 1997.

4. S. Alford and E. Schwartz. Presynaptic inhibition. In Encyclopedia of Neuroscience, pages 1001–1006. Academic Press.

5. D. J. Amit and N. Brunel. Model of global spontaneous activity and local structured activity during delay periods in the cerebral cortex. Cerebral cortex (New York, NY: 1991), 7(3):237–252, 1997.

6. E. L. Bienenstock, L. N. Cooper, and P. W. Munro. Theory for the development of neuron selectivity: orientation specificity and binocular interaction in visual cortex. The Journal of Neuroscience, 2(1):32–48.

7. S. Blomfield. Arithmetical operations performed by nerve cells. Brain research, 69(1):115–124, 1974.

8. N. Bowery, A. Hudson, and G. Price. Gabaa and gabab receptor site distribution in the rat central nervous system. Neuroscience, 20(2):365–383, 1987.

9. T. Branco and K. Staras. The probability of neurotransmitter release: variability and feedback control at single synapses. Nature Reviews Neuroscience, 10(5):373–383, 2009.

10. T. Branco, K. Staras, K. J. Darcy, and Y. Goda. Local dendritic activity sets release probability at hippocampal synapses. Neuron, 59(3):475–485, 2008.

11. N. Brunel. Dynamics of sparsely connected networks of excitatory and inhibitory spiking neurons. Journal of Computational Neuroscience, 8(3):183–208, 2000.

12. F. S. Chance, L. F. Abbott, and A. D. Reyes. Gain modulation from background synaptic input. Neuron, 35(4):773–782, 2002.

13. C. Chen and W. G. Regehr. Presynaptic modulation of the retinogeniculate synapse. The Journal of Neuroscience, 23(8):3130–3135, 2003.

14. A. J. Delaney, J. W. Crane, N. M. Holmes, J. Fam, and R. F. Westbrook. Baclofen acts in the central amygdala to reduce synaptic transmission and impair context fear conditioning. Scientific Reports, 8(1):9908, 2018.

15. N. S. Desai, L. C. Rutherford, and G. G. Turrigiano. Plasticity in the intrinsic excitability of cortical pyramidal neurons. Nature neuroscience, 2(6):515, 1999.

16. J. S. Dittman and W. G. Regehr. Mechanism and kinetics of heterosynaptic depression at a cerebellar synapse. Journal of Neuroscience, 17(23):9048–9059, 1997.

17. H. S. Engelman and A. B. MacDermott. Presynaptic ionotropic receptors and control of transmitter release. Nature Reviews Neuroscience, 5(2):135, 2004.

18. A. J. P. Fink, K. R. Croce, Z. J. Huang, L. F. Abbott, T. M. Jessell, and E. Azim. Presynaptic inhibition of spinal sensory feedback ensures smooth movement. Nature, 509(7498):43–48, 2014.

19. J. Gjorgjieva, C. Clopath, J. Audet, and J.-P. Pfister. A triplet spike-timing–dependent plasticity model generalizes the bienenstock–cooper–munro rule to higher-order spatiotemporal correlations. Proceedings of the National Academy of Sciences, 108(48):19383–19388, 2011.

20. D. O. Hebb. The Organisation of Behaviour: A Neuropsychological Theory. Wiley, 1949.

21. G. R. Holt and C. Koch. Shunting inhibition does not have a divisive effect on firing rates. Neural computation, 9(5):1001–1013, 1997.

22. J. S. Isaacson, J. M. Solis, and R. A. Nicoll. Local and diffuse synaptic actions of GABA in the hippocampus. Neuron, 10(2):165–175, 1993.

23. E. M. Izhikevich. Dynamical systems in neuroscience. MIT press, 2007.

24. T. Keck, T. Toyoizumi, L. Chen, B. Doiron, D. E. Feldman, K. Fox, W. Gerstner, P. G. Haydon, M. Hübener, H.-K. Lee, et al. Integrating hebbian and homeostatic plasticity: the current state of the field and future research directions. Philosophical Transactions of the Royal Society B: Biological Sciences, 372(1715):20160158, 2017.

25. T. Laviv, I. Riven, I. Dolev, I. Vertkin, B. Balana, P. A. Slesinger, and I. Slutsky. Basal GABA regulates GABABR conformation and release probability at single hippocampal synapses. Neuron, 67(2):253–267, 2010.

26. J. Li, E. Park, L. R. Zhong, and L. Chen. Homeostatic synaptic plasticity as a metaplasticity mechanism—a molecular and cellular perspective. Current opinion in neurobiology, 54:44–53, 2019.

27. A. B. MacDermott, L. W. Role, and S. A. Siegelbaum. Presynaptic ionotropic receptors and the control of transmitter release. Annual review of neuroscience, 22(1):443–485, 1999.

28. H. Markram and M. Tsodyks. Redistribution of synaptic efficacy between neocortical pyramidal neurons. Nature, 382(6594):807, 1996.

29. H. Markram, Y. Wang, and M. Tsodyks. Differential signaling via the same axon of neocortical pyramidal neurons. Proceedings of the National Academy of Sciences, 95(9):5323–5328, 1998.

30. R. J. Miller. Presynaptic receptors. Annual review of pharmacology and toxicology, 38(1):201–227, 1998.

31. S. J. Mitchell and R. A. Silver. Shunting inhibition modulates neuronal gain during synaptic excitation. Neuron, 38(3):433–445, 2003.

32. B. J. Molyneaux and M. E. Hasselmo. GABAB presynaptic inhibition has an in vivo time constant sufficiently rapid to allow modulation at theta frequency. Journal of Neurophysiology, 87(3):1196–1205, 2002.

33. R. Nicoll and B. Alger. Presynaptic inhibition: transmitter and ionic mechanisms. In International review of neurobiology, volume 21, pages 217–258. Elsevier, 1979.

34. H. Ohta and Y. P. Gunji. Recurrent neural network architecture with pre-synaptic inhibition for incremental learning. Neural Networks, 19(8):1106–1119, 2006.

35. S. R. Olsen and R. I. Wilson. Lateral presynaptic inhibition mediates gain control in an olfactory circuit. Nature, 452(7190):956–960, 2008.

36. A. Orts-Del’Immagine and J. R. Pugh. Activity-dependent plasticity of presynaptic gabab receptors at parallel fiber synapses. Synapse, 72(5):e22027, 2018.

37. L. Overstreet-Wadiche and C. J. McBain. Neurogliaform cells in cortical circuits. Nature Reviews Neuroscience, 16(8):458, 2015.

38. J.-P. Pfister. Triplets of spikes in a model of spike timing-dependent plasticity. Journal of Neuroscience, 26(38):9673–9682, 2006.

39. S. A. Prescott and Y. De Koninck. Gain control of firing rate by shunting inhibition: roles of synaptic noise and dendritic saturation. Proceedings of the National Academy of Sciences, 100(4):2076–2081, 2003.

40. J. S. Rothman, L. Cathala, V. Steuber, and R. A. Silver. Synaptic depression enables neuronal gain control. Nature, 457(7232):1015, 2009.

41. J. Serrats, M. O. Cunningham, and C. H. Davies. Gabab receptor modulation – to b or not to be b a pro-cognitive medicine? Current opinion in pharmacology, 35:125–132, 2017.

42. H. Shaban, Y. Humeau, C. Herry, G. Cassasus, R. Shigemoto, S. Ciocchi, S. Barbieri, H. v. d. Putten, K. Kaupmann, B. Bettler, and A. Lüthi. Generalization of amygdala LTP and conditioned fear in the absence of presynaptic inhibition. Nature Neuroscience, 9(8):1028–1035, 2006.

43. P. J. Sjöström, G. G. Turrigiano, and S. B. Nelson. Rate, timing, and cooperativity jointly determine cortical synaptic plasticity. Neuron, 32(6):1149–1164, 2001.

44. E. Slomowitz, B. Styr, I. Vertkin, H. Milshtein-Parush, I. Nelken, M. Slutsky, and I. Slutsky. Interplay between population firing stability and single neuron dynamics in hippocampal networks. eLife, 4:e04378, 2015.

45. H. Sprekeler. Functional consequences of inhibitory plasticity: homeostasis, the excitation-inhibition balance and beyond. Current opinion in neurobiology, 43:198–203, 2017.

46. M. Stimberg, D. F. Goodman, V. Benichoux, and R. Brette. Equation-oriented specification of neural models for simulations. Frontiers in neuroinformatics, 8:6, 2014.

47. M. Tachibana and A. Kaneko. Retinal bipolar cells receive negative feedback input from gabaergic amacrine cells. Visual neuroscience, 1(3):297–305, 1988.

48. S. M. Thompson, M. Capogna, and M. Scanziani. Presynaptic inhibition in the hippocampus. Trends in Neurosciences, 16(6):222–227, 1993.

49. M. Tsodyks, K. Pawelzik, and H. Markram. Neural networks with dynamic synapses. Neural computation, 10(4):821–835, 1998.

50. M. V. Tsodyks and H. Markram. The neural code between neocortical pyramidal neurons depends on neurotransmitter release probability. Proceedings of the national academy of sciences, 94(2):719–723, 1997.

51. G. G. Turrigiano, K. R. Leslie, N. S. Desai, L. C. Rutherford, and S. B. Nelson. Activity-dependent scaling of quantal amplitude in neocortical neurons. Nature, 391(6670):892–896.

52. G. G. Turrigiano and S. B. Nelson. Homeostatic plasticity in the developing nervous system. Nature Reviews Neuroscience, 5(2):97, 2004.

53. J. Urban-Ciecko, E. Fanselow, and A. Barth. Neocortical somatostatin neurons reversibly silence excitatory transmission via GABAb receptors. Current Biology, 25(6):722–731, 2015.

54. R. Vigot, S. Barbieri, H. Bräuner-Osborne, R. Turecek, R. Shigemoto, Y.-P. Zhang, R. Luján, L. H. Jacobson, B. Biermann, J.-M. Fritschy, C.-M. Vacher, M. Müller, G. Sansig, N. Guetg, J. F. Cryan, K. Kaupmann, M. Gassmann, T. G. Oertner, and B. Bettler. Differential compartmentalization and distinct functions of GABAB receptor variants. Neuron, 50(4):589–601, 2006.

55. T. P. Vogels and L. F. Abbott. Signal propagation and logic gating in networks of integrate-and-fire neurons. Journal of neuroscience, 25(46):10786–10795, 2005.

56. T. P. Vogels, H. Sprekeler, F. Zenke, C. Clopath, and W. Gerstner. Inhibitory plasticity balances excitation and inhibition in sensory pathways and memory networks. Science, 334(6062):1569–1573, 2011.

57. L. Wang, M. Kloc, E. Maher, A. Erisir, and A. Maffei. Presynaptic gabaa receptors modulate thalamocortical inputs in layer 4 of rat v1. Cerebral Cortex, 29(3):921–936, 2018.

58. L.-G. Wu and P. Saggau. Presynaptic inhibition of elicited neurotransmitter release. Trends in Neurosciences, 20(5):204–212, 1997.

59. F. Zenke and W. Gerstner. Hebbian plasticity requires compensatory processes on multiple timescales. Philosophical Transactions of the Royal Society B: Biological Sciences, 372(1715):20160259, 2017.

60. F. Zenke, G. Hennequin, and W. Gerstner. Synaptic plasticity in neural networks needs homeostasis with a fast rate detector. PLoS Computational Biology, 9(11).

61. D. Zhang, S. Wu, and M. J. Rasch. Circuit motifs for contrast-adaptive differentiation in early sensory systems: the role of presynaptic inhibition and short-term plasticity. PloS one, 10(2):e0118125, 2015.

